# *GNL3* is an evolutionarily-conserved stem cell gene influencing cell proliferation, animal growth, and regeneration in the hydrozoan *Hydractinia*

**DOI:** 10.1101/2022.04.24.489303

**Authors:** Gonzalo Quiroga-Artigas, Danielle de Jong, Christine E. Schnitzler

## Abstract

*Nucleostemin* (*NS*) is a vertebrate gene preferentially expressed in stem and cancer cells, which acts to regulate cell cycle progression, genome stability, and ribosome biogenesis. *NS* and its paralogous gene, *GNL3-like* (*GNL3L*), arose in the vertebrate clade after a duplication event from their orthologous gene, *G protein Nucleolar 3* (*GNL3*). Research on invertebrate *GNL3*, however, has been limited. To gain a greater understanding of the evolution and functions of the *GNL3* gene, we have performed studies in the hydrozoan cnidarian *Hydractinia symbiolongicarpus*, a colonial hydroid that continuously generates pluripotent stem cells throughout its life cycle and presents impressive regenerative abilities. We show that *Hydractinia GNL3* is expressed in stem and germline cells. The knockdown of *GNL3* reduces the number of mitotic and S-phase cells in *Hydractinia* larvae of different ages. Genome editing of *Hydractinia GNL3* via CRISPR/Cas9 resulted in colonies with reduced growth rates, polyps with impaired regeneration capabilities, gonadal morphological defects, and low sperm motility. Collectively, our study shows that *GNL3* is an evolutionarily-conserved stem cell and germline marker involved in cell proliferation, animal growth, regeneration, and sexual reproduction in *Hydractinia*, and sheds new light into the evolution of GNL3 and of stem cell systems.

## 1. Introduction

The study of stem cell biology has a long history, and is at the forefront of research in regenerative medicine. While much of our current understanding of stem cells is based on studies performed on a small number of model organisms (e.g., model vertebrates, *C. elegans*, and *D. melanogaster*), a complete understanding of the molecular basis and evolution of stem cell systems can only be gained when looking outside the conventional experimental models [1–3].

Cnidarians have proven to be excellent experimental models to study stem cell biology and evolution, due to their morphological simplicity, cellular plasticity, outstanding regenerative capabilities, long lifespan, and their key position in the phylogenetic tree as sister group to bilaterian animals [4, 5]. Within the Cnidaria, hydrozoans have been widely used to gain an understanding of stem cells and regeneration processes [4–6]. Notably, hydrozoans possess a population of undifferentiated stem/progenitor cells, which are generally proliferative and migratory, named interstitial cells (i-cells) [4–8].

The hydrozoan *Hydractinia symbiolongicarpus* (hereafter *Hydractinia*) is a marine, dioecious, colonial animal that lives on the surface of hermit crab shells in the wild (figure 1A). Its transparency, small size, and ease of manipulation and rearing, as well as a wide availability of molecular and genetic approaches, make *Hydractinia* a highly attractive research organism for experimental biology [9]. I-cells in *Hydractinia* derive from the embryonic endoderm, are found both in larval and adult stages (figure 1B), and are capable of giving rise to all somatic lineages as well as to germ cells [7,8,10]. It is currently unknown whether *Hydractinia* i-cells exist as a uniform population of pluripotent stem cells or as heterogeneous sub-populations of undifferentiated cells with mixed potencies. To date, in-depth functional analyses of *Hydractinia* i-cell genes are scarce [10–13], and none has been functionally characterized throughout the *Hydractinia* life cycle. Performing reverse genetics on *Hydractinia* i-cell genes and studying their function at different life cycle stages will shed light on the molecular basis and evolution of stem cell systems.

**Figure 1.**
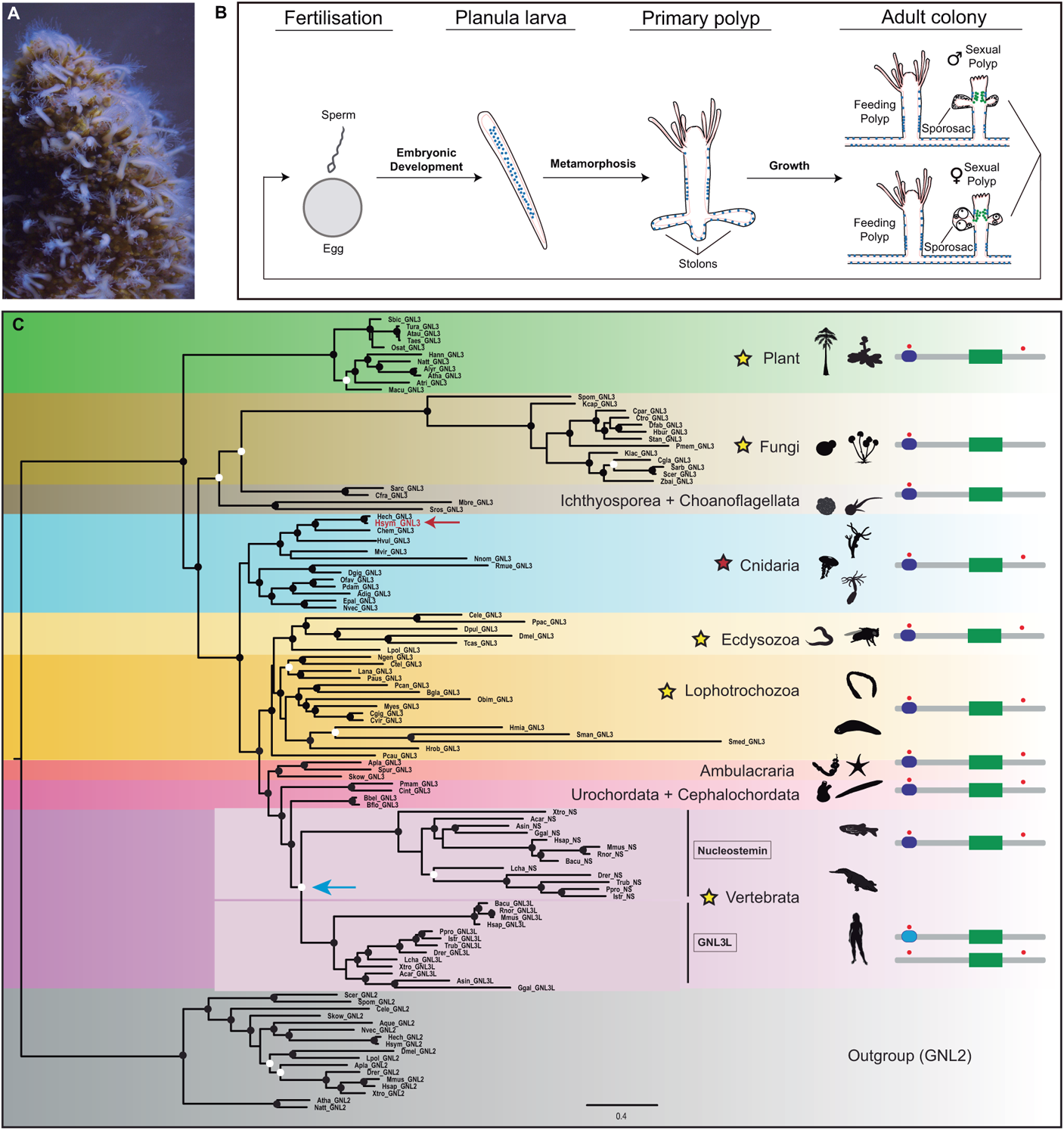
Molecular phylogeny and domain analysis of GNL3. (A) Photo of a *Hydractinia* colony on top of hermit crab shell. (B) Schematic of *Hydractinia* life cycle. Ectodermal and endodermal epithelial layers are depicted in black and pink, respectively. I-cells are represented by blue dots and germ cells by green dots. (C) Maximum likelihood (ML) phylogenetic analysis of GNL3 family of proteins (RAxML, LG + I + gamma + F; see electronic supplementary material, table S1 for full species names and accession numbers, and file S2, and S3 for input sequences and sequence alignment, respectively). Tree was rooted with GNL2 protein sequences. ML bootstrap support values (500 replicates) are shown as black circles on branch tips when support is higher than 70% and as white circles when support is higher than 50% but lower than 70%. Scale bar shows estimated number of substitutions per site. The subject of this study, *H. symbiolongicarpus* GNL3, is shown in red font and indicated by a red arrow. A putative duplication event in the vertebrate lineage, resulting in Nucleostemin and GNL3L paralogs is indicated by a blue arrow. To the right of the tree, yellow stars indicate phyla/supergroups where at least one study has been conducted, including this study. On the far right, schematics of the domain structure of representative GNL3 proteins are shown. Grey bar indicates entire length of protein, dark blue rounded rectangle represents presence of a GN3L_Grn1 putative GTPase domain, and green rectangle represents presence of a MMR1_HSR1 50S ribosome-binding GTPase domain. Light blue rounded rectangle in the domain structure of vertebrate GNL3L indicates a derived GN3L_Grn1 putative GTPase domain, while other vertebrate GNL3L proteins lack the domain entirely. Red circles indicate the presence of at least one nuclear localisation signal (NLS) in regions close to the N-terminus and C-terminus. Schematics are not to scale. Animal, plant, and fungi silhouettes were taken from http://phylopic.org.

G protein Nucleolar 3 (GNL3) is a nucleolar protein that belongs to the YlqF/YawG GTPase family, which is characterized by the presence of an MMR1_HSR1 domain of five circularly permuted GTP-binding motifs [14]. The *GNL3* gene is present as a single copy in non-vertebrate eukaryotes. A presumptive duplication event in the vertebrate clade gave rise to two paralogs, *nucleostemin* (*NS*) and *GNL3-like* (*GNL3L*) [14, 15] (figure 1C).

The *NS* gene is highly expressed in cells capable of continuous proliferation such as different types of stem cells, including embryonic and adult neuronal stem cells, primordial germ cells, and tumour cells [14, 16]. Moreover, NS contributes to biological processes such as embryogenesis, tissue regeneration, and cancer development [17], making *NS* a target gene for stem cell and cancer research, as well as for regenerative and reproductive medicine. NS is capable of shuttling between the nucleolus and the nucleoplasm depending on its GTP binding state [18], and interacts directly or indirectly with a large number of proteins [14, 15]. NS is involved in a variety of cellular functions such as cell cycle regulation and self-renewal [14,15,17,19], genome integrity [20, 21], and ribosomal biogenesis [22–24], and can act as a reprogramming factor [19], altogether making NS a multifunctional protein essential for stem cell regulation.

The vertebrate *NS* paralog, *GNL3L*, has been less studied overall. Unlike *NS*, *GNL3L* shows lower expression in stem cells and higher expression in differentiated tissues, although it is found upregulated in some cancers [14]. GNL3L is also able to shuttle between the nucleolus and the nucleoplasm, with a shorter nucleolar residence than NS [18]. GNL3L has been shown to regulate cell cycle transitions [25, 26], to negatively regulate telomere length [27], to modulate the transcriptional levels of oestrogen-related receptors [28], and to mediate pre-ribosomal RNA processing [24].

Invertebrate *GNL3* is a term that is used to refer to the *GNL3* gene from invertebrate animals, but also from organisms of other kingdoms like fungi and plants [17]. *GNL3* was recently identified as one of 195 genes found to have enriched expression in multipotent stem cells across several invertebrate animals, including the sponge *S. lacustris*, the cnidarian *H. vulgaris*, the planarian *S. mediterranea*, and the schistosome *S. mansoni* [29]. Only a limited number of functional studies have been performed on invertebrate *GNL3*, however, and these have mostly been focused on typical model organisms. In *S. cerevisiae*, GNL3 (Nug1p) is necessary for its viability and for 60S ribosomal subunit export [30], while in *S. pombe*, GNL3 (Grn1p) is required for growth, pre-ribosomal RNA processing, and nucleolar export of pre-ribosomal complexes [31]. In *D. melanogaster*, GNL3 (NS1) is crucial for larval and pupal development, for the nucleolar release of large ribosomal subunits, and for the maintenance of larval midgut precursor cells [32]. In *C. elegans*, GNL3 (NST-1) is needed for larval growth, germline stem cell proliferation, and is involved in ribosome biogenesis [33]. In the planarian *S. mediterranea*, *GNL3* (*Nucleostemin*) is needed for complete head and tail regeneration [34]. In *A. thaliana*, *GNL3* (*nsn1*) is highly expressed in embryos, in shoot and floral apical undifferentiated (meristematic) cells, and in developing leaves and flowers [35, 36], and is required for the maintenance of inflorescence meristem identity, embryonic development, cell proliferation, plant growth, fertility, and ribosome biogenesis [35–38]. Importantly, despite this evidence of *GNL3* stem cell and germ cell expression and function in these organisms, the current evolutionary paradigm that has been proposed is that vertebrate *GNL3L* is the direct descendant of invertebrate *GNL3*, and that *NS* arose as a novel gene with new functions during vertebrate evolution [17, 24].

Here, we identify and characterize *GNL3* in the hydrozoan cnidarian *Hydractinia* as an evolutionarily-conserved stem cell gene. We show that *GNL3* is expressed in i-cells throughout the *Hydractinia* life cycle, as well as in the germline. Gene functional analyses demonstrate that *GNL3* disruption affects cell proliferation, colony growth, polyp head regeneration, and sperm motility. By performing a broad cross-kingdom molecular phylogeny and domain analysis of GNL3 amino acid sequences, we show that a domain named GN3L_Grn1, located near the N-terminus, is shared between invertebrate GNL3 and vertebrate NS, but is mostly absent in the NS paralog GNL3L. We discuss how our findings contrast with the current paradigm of *GNL3* gene evolution, and propose an alternative evolutionary scenario. Our study helps to highlight the importance of *GNL3* in cancer and stem cell research, as well as the significance of studying diverse animals, and of performing wide phylogenetic comparisons, to better understand gene evolution.

## 2. Results

### 2.1. GNL3 molecular phylogeny and domain analysis

Homologues belonging to the YlqF/YawG GTPase family (GNL3, GNL2, LSG1, MTG1) were identified from BLAST searches of the *Hydractinia* genome and transcriptome databases. We could not find a *Hydractinia* ortholog of GNL1. Protein cluster map analyses demonstrate the high sequence conservation of *Hydractinia* GNL3, GNL2, and LSG1 to other animal orthologues of the same gene subfamilies (electronic supplementary material, figure S1).

Detailed phylogenetic analyses of GNL3 amino acid sequences encompassing different kingdoms, superphyla, phyla, and subphyla were performed using maximum likelihood (figure 1C). The GNL2 subfamily of proteins was used as the outgroup. In preliminary analyses we included sequences from additional non-bilaterian groups (ctenophore, sponge, placozoa) but these sequences tended to have a shifting placement within the trees with low levels of support. Since their position in the tree was ambiguous, and thus not informative, we decided to exclude them from further analyses. A single copy of GNL3 is present in plants, fungi, protists, and invertebrate animals. For the most part, the GNL3 tree we constructed is congruent with our current understanding of the species tree for the clades included in the analysis. This congruence is typical when there is a single copy of the gene for every species included without duplications or losses. The *Hydractinia* GNL3 sequence falls within the hydrozoan cnidarian group, as expected. Our tree supports the putative duplication event in the vertebrate clade which led to the formation of the paralogs GNL3L and NS, and all vertebrates we surveyed had both paralogs which group in their own clades (figure 1C).

While focusing on protein sequence comparisons, *Hydractinia* GNL3, as well as other invertebrate GNL3 sequences, have a slightly higher overall pairwise similarity to vertebrate GNL3L than to vertebrate NS (electronic supplementary material, figure S1). Specifically, *Hydractinia* GNL3 has a 34.7% pairwise identity with human NS and a 38.4% pairwise identity with human GNL3L. Sequence identity levels, however, do not necessarily provide a full picture of how proteins evolve and need to be complemented by the study of the presence/absence of functional protein domains to better infer sequence evolution. To assess this, we performed PFAM and MotifScan analyses and compared the domain structure of NS/GNL3L from several clades with *Hydractinia* GNL3 and other invertebrate GNL3 protein sequences. All amino acid sequences displayed a MMR1-HSR1 domain, expected in the YlqF/YawG GTPase family of proteins, and most of them had one to several nuclear localization signals (NLS) at the N-terminus, and sometimes also at the C-terminus, of their sequences (figure 1C). Importantly, invertebrate GNL3 proteins, including *Hydractinia* GNL3, share a GN3L_Grn1 domain at the N-terminus of the protein sequence with vertebrate NS, however this domain is often not present in vertebrate GNL3L, and when present, it is detected with low e-values by PFAM, indicating poor domain identity (figure 1C). These analyses strongly suggest that vertebrate GNL3L is more evolutionarily derived than vertebrate NS, and that there may be more functional protein conservation between NS and invertebrate GNL3 than between GNL3L and invertebrate GNL3.

### 2.2. The *Hydractinia GNL3* gene is expressed in larval and adult i-cells, germ cells, oocytes, and spermatogonia

In *Hydractinia* larvae, i-cells are located in the endoderm and migrate through the mesoglea to the larval ectoderm during metamorphosis [7]. Labelling with 5-ethynyl-2’-deoxyuridine (EdU), a thymidine analogue that is incorporated into the DNA of cells during S-phase, confirmed that the majority of proliferating cells were present in the larval endoderm (electronic supplementary material, figure S2A-B). Whole mount *in situ* hybridisation (ISH) of *Hydractinia* 1 dpf (days post-fertilisation), 2 dpf, and 3 dpf larvae revealed *GNL3* expression in a population of endodermal cells (figure 2A-C). Double fluorescent ISH on 2 dpf larvae using probes for *GNL3* and *Piwi1* (an i-cell marker), or *GNL3* and *PCNA* (a cell proliferation marker), revealed that both these genes were co-expressed with *GNL3* in a subset of cells within the larval endoderm (figure 2D-F). Detection of *GNL3* probe in parallel with EdU also revealed co-labelling of these two markers in a subset of cells (figure 2G). These findings indicate that *GNL3* is expressed in i-cells and, more generally, in proliferating cells within the endoderm of *Hydractinia* larvae.

**Figure 2.**
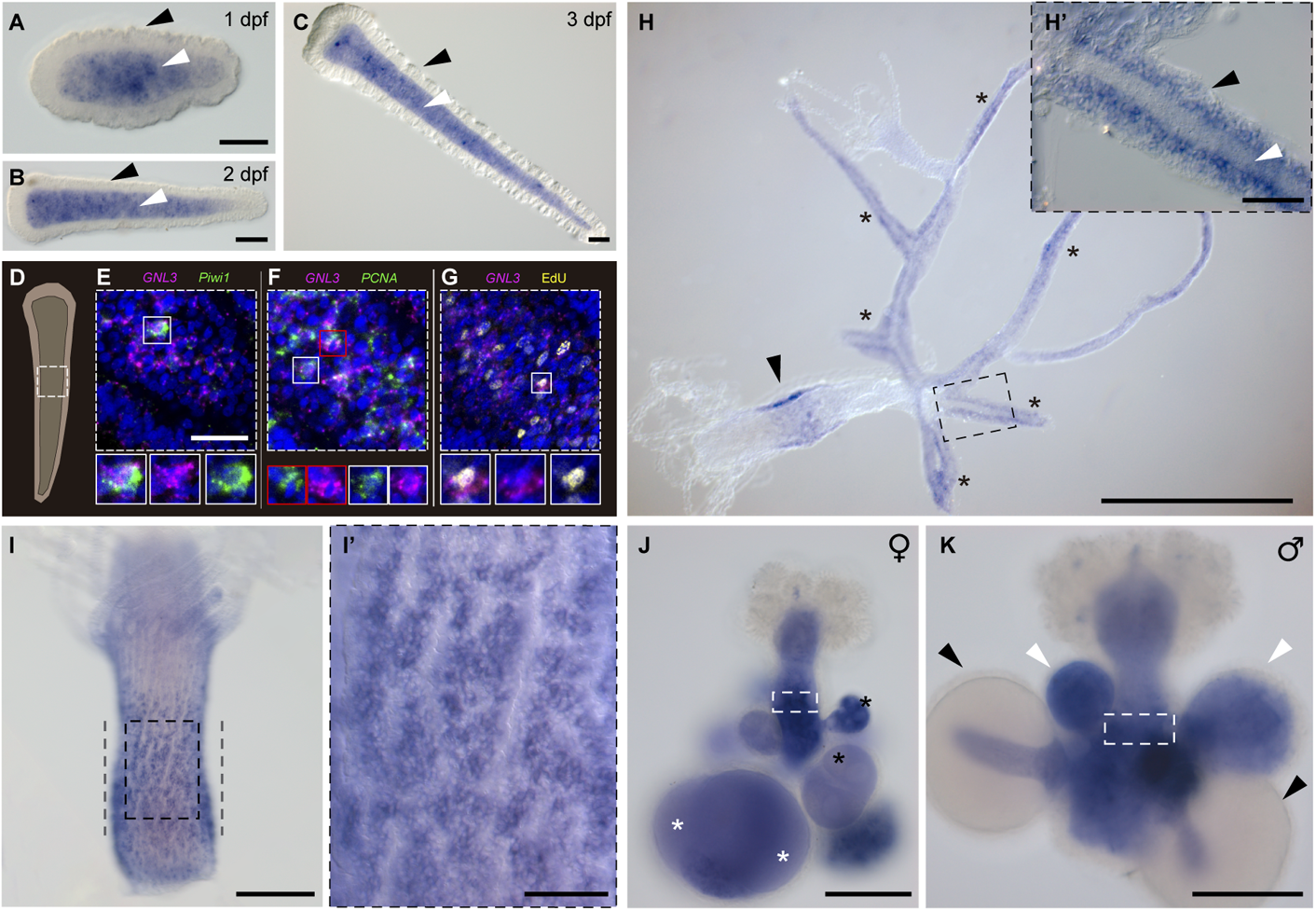
Whole mount ISH of *GNL3* in *Hydractinia*. (A-C) *GNL3* expression in 1 dpf, 2 dpf, and 3 dpf *Hydractinia* larvae is detected in the larval endoderm (white arrowheads) but not in the larval ectoderm (black arrowheads). (D) Schematic of a 2 dpf larva. White square defines approximately the larval region imaged in E-G. (E-F) Double fluorescent ISH showing co-expression of *GNL3* (magenta) and *Piwi1* or *PCNA* (green) in a subset of larval endodermal cells. (G) Fluorescent ISH of *GNL3* co-stained with EdU (yellow) showing some larval endodermal cells labelled with both markers. For E-G, nuclei are in blue, and examples of co-expression/co-staining are indicated by white and red boxes. (H) *GNL3* expression in a young colony with expression in the primary polyp (arrowhead) and stolons (asterisks). (H’) Magnification of region outlined in H showing *GNL3* expression in the stolonal epidermis (black arrowhead) but not in the gastrodermis (white arrowhead). (I) *GNL3* expression in the i-cell band-like area (defined by grey dashed lines) of an adult feeding polyp. (I’) Magnification of region outlined in I showing *GNL3* expression in cell clusters. (J) *GNL3* expression in the germinal zone (white rectangle), in growing oocytes (black asterisks), and in fully-grown oocytes (white asterisks) of female sexual polyps. (K) *GNL3* expression in the germinal zone (white rectangle), and in spermatogonia within small and medium-sized sporosacs (white arrowheads) of male sexual polyps. Note the absence of *GNL3* expression in mature sperm within large sporosacs (black arrowheads). ISH = *in situ* hybridisation; dpf = days post-fertilisation. Scale bars: 50 μm in A-C, H’, I’; 25 μm in E-G; 500 μm in H; 100 μm in I-K.

We could also observe *GNL3* expression in stolons, within the interstitial region of the epidermis known to be inhabited by i-cells (figure 2H-H’) [39], and in the interstitial region of the epidermis in a band-like area in primary polyps (figure 2H) and adult feeding polyps (figure 2I-I’), which is also known to host i-cells [11]. The *GNL3* expression pattern is reminiscent of, but more restricted than the distribution of EdU^+^ cells in primary polyps and adult feeding polyps (electronic supplementary material, figure S2C-D). These results strongly suggest that *GNL3* is expressed in feeding polyp and stolonal i-cells of juvenile and adult *Hydractinia* colonies.

In sexual polyps, we detected *GNL3* expression in the germinal zone bearing early germ cells [10] of both female and male sexual polyps (figure 2J-K; electronic supplementary material, figure S3A-B). We also found *GNL3* expressed in both growing and fully-grown oocytes in all sporosacs of female sexual polyps (figure 2J), whereas in male sexual polyps, *GNL3* probe stained spermatogonia inside small and medium-sized sporosacs, but not the mature sperm present in larger sporosacs (figure 2K). When combining EdU labelling with *GNL3* fluorescent ISH in male sexual polyps, we observed complete co-localization in spermatogonia within small and medium-sized sporosacs, but the complete absence of both markers in larger sporosacs (electronic supplementary material, figure S3A). These results illustrate that *GNL3* is expressed in germ cells, oocytes and proliferating spermatogonia of female and male sexual polyps.

### 2.3. GNL3 knockdown reduces the number of proliferating and mitotic cells in larvae

To better understand the functions of GNL3 in larvae, we knocked down the transcript of *Hydractinia GNL3* via short hairpin RNA (shRNA) electroporation of one-cell stage embryos, then studied different cellular markers in larvae of different ages following knockdown (KD). We designed and synthesised a single shRNA targeting our gene of interest, and electroporated *Hydractinia* fertilised eggs at a concentration of 1500 ng/μl (see methods). As a negative control for these experiments, we used a scrambled sequence of the shRNA used for *GNL3* knockdown, and verified that it did not target any gene in the *Hydractinia* genome using a BLAST homology search. We obtained strikingly reduced levels of *GNL3* mRNA, as shown by qPCR, in larval samples from different ages (figure 3A), while observing equivalent survivorship levels between control and *GNL3* KD larvae (76.9% ± 5.6% and 76.1% ± 3.3%, respectively; n=7 independent experiments).

**Figure 3.**
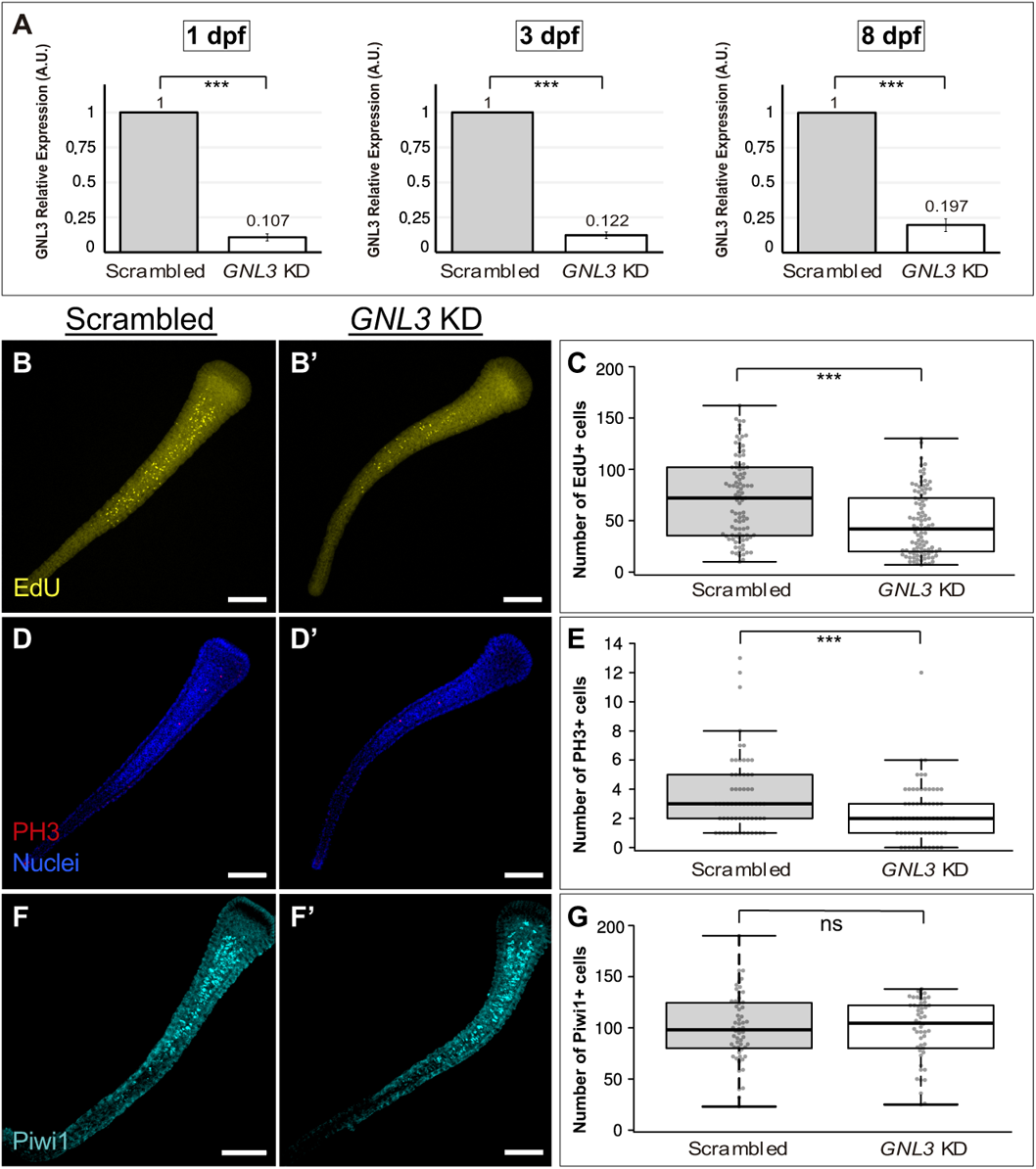
*GNL3* knockdown significantly reduces the number of EdU^+^ and PH3^+^ cells without affecting the number of Piwi1^+^ cells in larvae. (A) RT-qPCR showing a significant decrease of *GNL3* transcript levels in *GNL3* shRNA-electroporated samples (*GNL3* KD) relative to scrambled shRNA controls at 1 dpf (left), 3 dpf (middle), and 8 dpf (right). Bar heights represent mean values of three independent experiments and error bars show standard deviations. A.U. arbitrary units. (B-B’, D-D’, F-F’) Representative images of 8 dpf larvae showing EdU^+^ cells (yellow, B-B’), PH3^+^ cells (red, D-D’), or Piwi1^+^ cells (cyan, F-F’) in scrambled and *GNL3* KD larvae. (C, E, G) Box plots showing the number of EdU^+^, PH3^+^, or Piwi1^+^ cells for scrambled and *GNL3* KD 8 dpf larvae. Centre lines show the medians; box limits indicate the 25th and 75th percentiles (first and third quartiles); whiskers extend 1.5 times the interquartile range from the 25th and 75th percentiles; each quantified sample is represented by a grey circle. In all cases, the full depth of the larvae was imaged and images shown were projected from confocal stacks. EdU quantifications were combined from 3 independent experiments (scrambled, n = 95; *GNL3* KD, n = 97), whereas PH3 and Piwi1 quantifications were combined from 2 independent experiments (scrambled n = 62; *GNL3* KD, n = 66 for PH3; scrambled, n = 56; *GNL3* KD, n = 50 for Piwi1). ns = non-significant, dpf = days post-fertilisation; *** = p-value≤0.01. All scale bars: 100 μm.

First, to assess cell proliferation and mitosis, we performed EdU labelling (an S-phase marker), and immunofluorescence analysis with a universal phosphorylated histone H3 (PH3) antibody (a mitotic marker), in *GNL3* KD larvae and scrambled controls. We observed an overall reduction in the number of both EdU^+^ and PH3^+^ cells in *GNL3* KD larvae at 2, 3, and 8 dpf, which was statistically significant in all cases but one (figure 3B-E; electronic supplementary material, figure S4).

To determine whether the lower number of S-phase and mitotic cells in *GNL3* KD larvae was simply due to a reduction in the total number of larval i-cells, we used a Piwi1 antibody, previously used as an i-cell marker for *Hydractinia* [10, 11]. There was no significant difference in the number of Piwi1^+^ cells between *GNL3* KD and scrambled control larval samples at 2 dpf and 8 dpf timepoints (figure 3F-G; electronic supplementary material, figure S4E-F), indicating that the overall number of i-cells was not affected upon *GNL3* knockdown. Altogether, our results show that the knockdown of *GNL3* decreases the number of S-phase and M-phase cells without affecting the overall number of i-cells in *Hydractinia* larvae.

### 2.4. *GNL3* knockdown does not induce spontaneous DNA damage or apoptosis in larvae and does not affect larval mature ribosomal RNA species

We sought to determine whether *Hydractinia GNL3* knockdown induced spontaneous DNA damage in larval proliferating cells. For this purpose, we used a commercial antibody against gamma-H2A.X (GH2A.X), a widely-used DNA damage marker labelling double-strand breaks [20,40,41]. To artificially induce DNA damage in proliferating cells, we incubated larvae in 20mM hydroxyurea (HU) for 3 hours prior to fixation. HU treatment depletes the endogenous nucleotide pool and consequently stalls the replication fork, generating DNA damage [20, 42]. We observed that HU-treated larvae presented a high number of GH2A.X^+^ cells (with a pattern reminiscent of that shown by EdU - electronic supplementary material, figure S2A-B). However, performing the same experiment on *GNL3* KD larvae in the absence of HU shows that *GNL3* KD does not induce spontaneous DNA damage in *Hydractinia* larvae of different ages, as shown by the almost complete absence of GH2A.X^+^ cells, which was equivalent to the scrambled control (electronic supplementary material, figure S5).

To study whether *GNL3* knockdown induced cell apoptosis in *Hydractinia* larvae, we used a TUNEL assay to label apoptotic cells. We observed that, whereas DNase I-treated larvae presented a vast number of TUNEL^+^ cells, *GNL3* KD larvae of different ages displayed an almost complete absence of TUNEL^+^ cells, comparable to the scrambled control (electronic supplementary material, figure S6). This result indicates that the downregulation of *GNL3* does not induce apoptosis in *Hydractinia* larvae.

We also aimed to assess whether mature ribosomal RNA (rRNA) species (18S and 28S) were affected when *GNL3* was downregulated in *Hydractinia* larvae. We extracted total RNA from scrambled control and *GNL3* KD larvae of different ages from three independent biological replicates, and used an Agilent 2100 Bioanalyzer Instrument to check the levels and ratios of mature 28S and 18S rRNA. We first observed that the levels of mature rRNA species were equivalent between conditions in all cases (electronic supplementary material, figure S7A). Next, by analysing the Bioanalyzer electropherogram outputs, we obtained non-significant differences (p-values>0.1) in the 28S / 18S rRNA ratios between conditions at all timepoints (for 2 dpf samples: scrambled 28S/18S = 2.03 ± 0.06; *GNL3* KD 28S/18S = 2.13 ± 0.21; for 3 dpf samples: scrambled 28S/18S = 1.87 ± 0.06; *GNL3* KD 28S/18S = 1.97 ± 0.15; for 8 dpf samples: scrambled 28S/18S = 2.00 ± 0.17; *GNL3* KD 28S/18S = 2.07 ± 0.25; electronic supplementary material, figure S7B-D). These results strongly suggest that the biosynthesis of mature rRNAs is not altered when *GNL3* is downregulated in *Hydractinia* larvae.

### 2.5. *GNL3* knockout hinders colony growth

Since the effects of a gene knockdown are transient, generally lasting for less than two weeks in *Hydractinia* [10, 43], to test the phenotypic effects of disrupting *GNL3* in *Hydractinia* colonies over a longer-term, we opted to create *GNL3* knockout (KO) lines using CRISPR/Cas9 technology. Genome editing via CRISPR-Cas9 has previously been successfully deployed in *Hydractinia*, achieving knockout lines for targeted genes [10, 44]. The *Hydractinia GNL3* gene consists of 14 exons and 13 introns. The overall GC content of the gene, including introns and exons, is 33.9% and that of the coding region is 40.2%. We designed three different CRISPR small guide RNAs (sgRNAs) targeting the 5^th^ and 6^th^ exons of the *GNL3* gene (figure 4A). We microinjected Cas9 protein and a mixture of the three sgRNAs into unfertilised eggs, followed by fertilisation (see methods). In parallel, we injected Cas9 protein without sgRNAs as our negative controls (Cas9-only). We designed primers flanking the targeted region of *GNL3* (electronic supplementary material, figure S8A) and performed PCRs using genomic DNA samples from individual larva developed from embryos injected with sgRNA/Cas9 complexes. We observed that all larvae presented *GNL3* gene editing, albeit in all cases mosaic (electronic supplementary material, figure S8B), indicating that our strategy was efficient in inducing mutations of our target gene. We utilised the inherent mosaicism of the F0 lines to our advantage, since a full knockout of *GNL3* might have provoked embryonic lethality, similar to what occurs upon *NS* depletion in mice [19, 45], deterring our ability to study its function.

**Figure 4.**
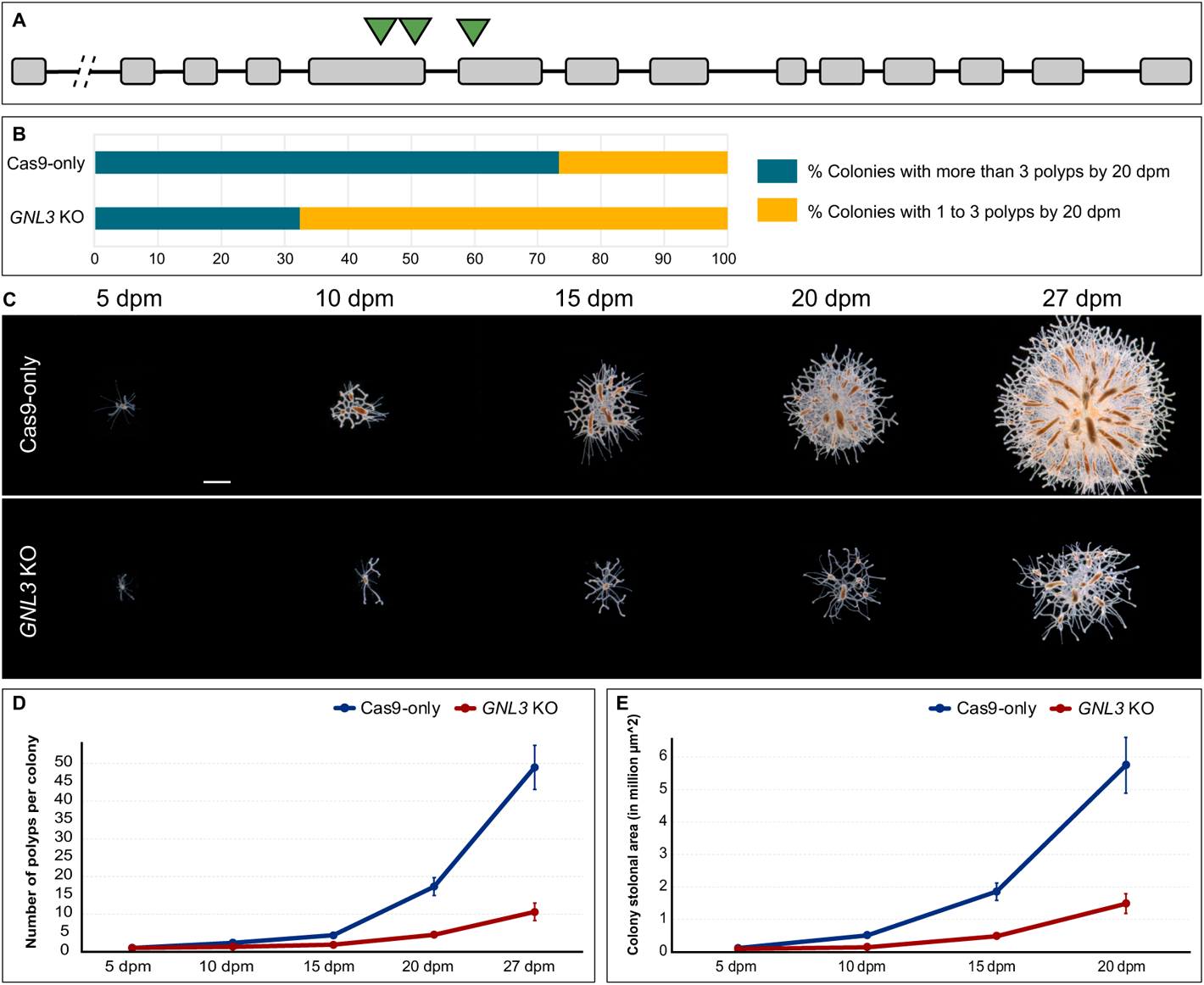
*GNL3* knockout slows colony growth. (A) *GNL3* gene structure depicting exons (grey boxes), introns (black lines) and target sites of sgRNAs (green triangles). Break between 1^st^ and 2^nd^ exon denotes several thousand base pairs of intronic region. (B) Graph depicting the percentage of colonies with more than 3 polyps by 20 dpm (green-blue) and the percentage of colonies with 1 to 3 polyps by 20 dpm (yellow) for Cas9-only and *GNL3* KO conditions. Fisher’s exact test showed a significant difference (p<0.01) between Cas9-only and *GNL3* KO samples. For Cas9-only, n = 162; for *GNL3* KO, n = 178. Data was combined from 2 independent experiments. (C) Representative images of Cas9-only and *GNL3* KO colonies at different timepoints post-metamorphosis. (D) Polyp number quantification at different timepoints in *Hydractinia* colonies for Cas9-only and *GNL3* KO conditions. (E) Stolonal area quantification at different timepoints in *Hydractinia* colonies for Cas9-only and *GNL3* KO conditions. In D-E, dots represent mean values and error bars show standard errors of the mean. Two-way ANOVA followed by post-hoc Bonferroni correction showed significant differences between conditions for both quantifications (electronic supplementary material, file S7). For Cas9-only, n = 44; for *GNL3* KO, n = 54. dpm = days post-metamorphosis. Scale bar = 1 mm.

A subset of embryos injected with sgRNA/Cas9 complexes and Cas9-only controls were reared to 21 dpf and subsequently labelled with EdU to assess changes in cell proliferation. In agreement with what we observed with younger larvae in our *GNL3* knockdown experiments (see above), we noted a significant reduction in the number of EdU^+^ cells in our *GNL3* KO larvae compared to Cas9-only controls (electronic supplementary material, figure S9). These results showed that we could phenocopy the effects of *GNL3* KD on cell proliferation in our *GNL3* KO larvae, which encouraged us to perform phenotypic analyses on adult colonies.

To study a potential *GNL3* KO phenotype related to colony growth, we metamorphosed 3dpf larvae injected with sgRNA/Cas9 complexes or Cas9-only, and obtained 92-98% metamorphosis success in both control and *GNL3* KO animals. This indicated that larval metamorphosis is not affected when the *GNL3* gene is disrupted. We then counted the percentage of colonies that presented more than 3 polyps by 20 dpm (days post-metamorphosis) and observed a much higher percentage of these colonies in the Cas9-only controls than in the *GNL3* KO animals (figure 4B). We performed PCR analyses using genomic DNA as template to determine the correlation between colony growth impairment and *GNL3* gene editing, and noted that 74% of the slow-growing colonies (colonies with 1 to 3 polyps by 20 dpm) from *GNL3* knockouts correlated to some level of *GNL3* gene editing. Genotyping by sequencing of multiple clones from three different sexually-mature colonies that presented slow growth (named 1.G2, 6.F3L, and 6.K2B) revealed multiple deletions in the *GNL3* gene, which produced non-sense and missense mutations when *in silico* translation on the mutant sequences was performed. In each, the wild type (WT) *GNL3* sequence was also detected, making these colonies mosaic *GNL3* mutants (electronic supplementary material, figure S10). Lastly, we conducted a time series experiment where we took images of control and *GNL3* KO colonies at different timepoints up to 27 dpm, and quantified the number of polyps present in each colony as well as the stolonal area (figure 4C). *GNL3* KO colonies showed a significant reduction in both growth parameters when compared to control colonies (figure 4D-E). Altogether, these results demonstrate that GNL3 plays a role in normal *Hydractinia* colony growth.

### 2.6. Polyp head regeneration is impaired in *GNL3* knockout colonies

Under optimal conditions, *Hydractinia* feeding polyps are capable of fully regenerating a functional head following decapitation within approximately 72 hours (Figure 5A; [11]). To test the potential involvement of GNL3 in *Hydractinia* polyp head regeneration, we first performed ISH using a *GNL3* probe in 24 hpd (hours post-decapitation) polyps and identified cells expressing *GNL3* in the i-cell band region and in the blastema (i.e., region of high cell proliferation; figure 5B-B’). ISH of 48 hpd polyps revealed *GNL3* expression in the band region and in the newly-formed tentacle buds (figure 5C-C’). The *GNL3* expression pattern in regenerating polyps suggests that *GNL3* might play an important role in polyp head regeneration.

**Figure 5.**
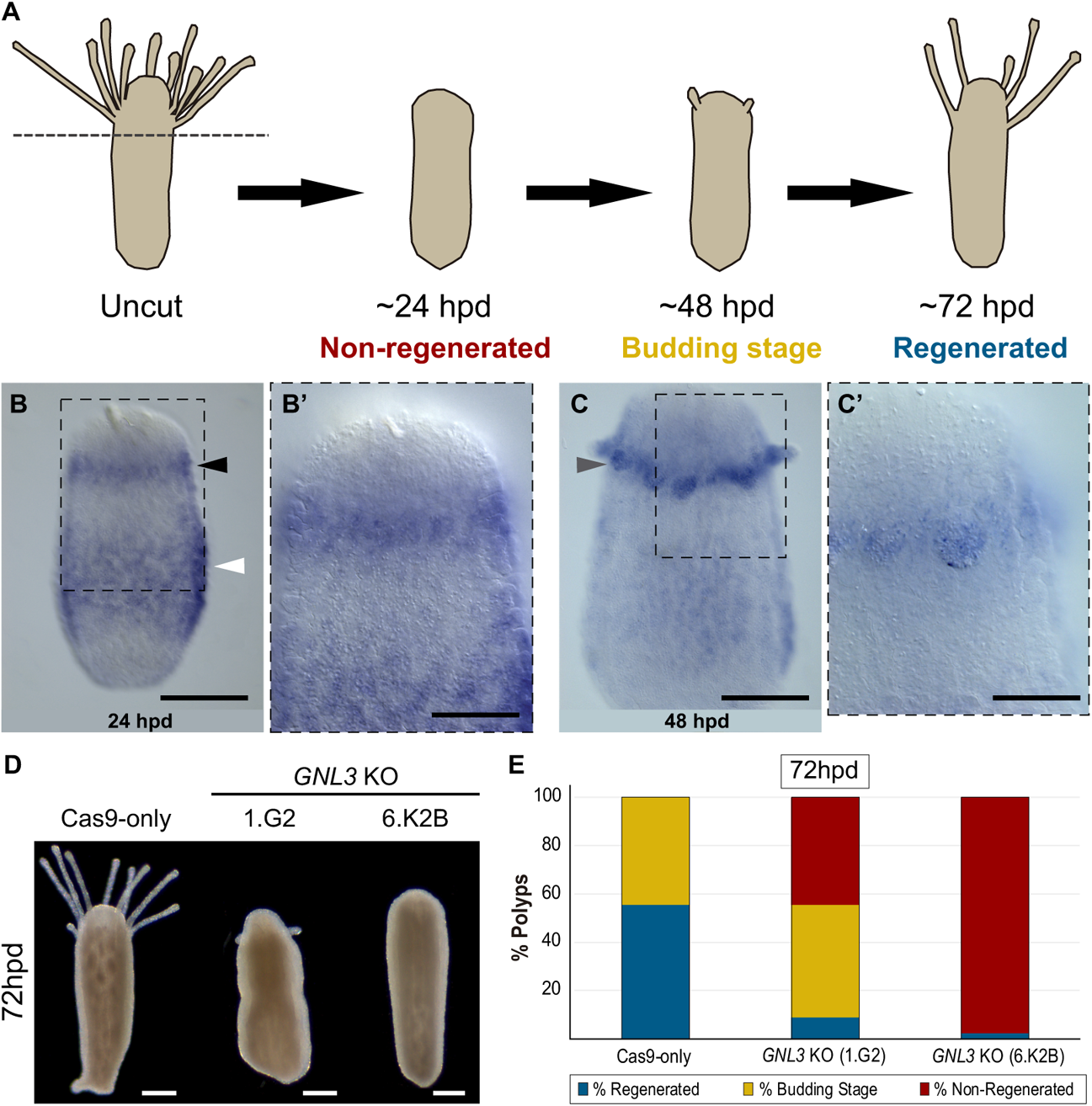
*GNL3* knockout impacts polyp head regeneration. (A) Schematics depicting *Hydractinia* feeding polyp head regeneration dynamics. Dotted grey line indicates position of decapitation. By 24 hpd polyps have not yet regenerated a head. By 48 hpd, new tentacle buds have formed. In about 72 hpd, polyps have regenerated a functional head with mouth and tentacles. (B) ISH detection of *GNL3* mRNA in the band area (white arrowhead) of a 24 hpd feeding polyp, as well as in the blastema region (black arrowhead). (B’) Magnification of region outlined in B. (C) ISH showing *GNL3* expression concentrated in the regenerating tentacle buds (grey arrowhead) of a 48 hpd feeding polyp. (C’) Magnification of region outlined in C. (D) Representative images of 72 hpd polyps from a Cas9-only control colony and from two *GNL3* KO colonies (1.G2 and 6.K2B). Note the slower or absent head regeneration in the *GNL3* KO colonies when compared to the control. (E) Stacked bar graph depicting the percentage of decapitated polyps that have regenerated (blue), are in a budding stage (yellow), or have not regenerated (red) by 72 hpd. In all cases, n = 45 decapitated polyps. Data was pooled from three independent experiments. hpd = hours post-decapitation. Scale bars: 100 μm in B, C; 50 μm in B’, C’; 200 μm in D.

To investigate if knockout of *GNL3* had an effect on *Hydractinia* polyp head regeneration, we dissected polyps from two *GNL3* mutant colonies (1.G2 and 6.K2B) as well as from age-matched Cas9-only control colonies, decapitated them, and assessed their ability to regenerate a head. Whereas over 50% of Cas9-only polyps had fully regenerated their heads by 72 hpd, with the remaining control polyps forming tentacle buds, the polyps from the two *GNL3* KO colonies had much lower rates of regeneration by 72 hpd (figure 5D-E). Strikingly, almost 100% of polyps from 6.K2B colony, the slowest-growing of all *GNL3* KO colonies we bred to sexual maturity, displayed complete regeneration failure (figure 5E). After 5-10 days, most 1.G2 polyps did regenerate a head, but 6.K2B polyps did not, and underwent aboral regeneration [11] after 2-3 weeks without polyp budding (not shown). These results indicate a requirement for GNL3 during *Hydractinia* polyp head regeneration.

### 2.7. *GNL3* knockout affects sexual polyp morphology and reduces sperm motility

To gain insight into the function of GNL3 in *Hydractinia* sexual polyps, we first examined the morphology and size of the sexual polyps in the genotyped *GNL3* KO colonies (1.G2, 6.F3L, and 6.K2B), which were all males. While no obvious differences in size and morphology could be observed between 1.G2 and age-matched Cas9-only control sexual polyps, we detected defects in the sporosac structure in 6.F3L sexual polyps, such as gastrodermis fusion into the epidermis, and an overall smaller size as well as underdeveloped oral regions in 6.K2B sexual polyps (electronic supplementary material, figure S10). These findings suggest that GNL3 is involved in the growth and morphogenesis of male sexual polyps in *Hydractinia*.

Since we detected *GNL3* expression in germ cells and in spermatogonia (electronic supplementary material, figure S3), we analysed the sperm of our genotyped *GNL3* KO colonies, aiming to identify potential problems. All colonies were able to spawn mature sperm upon light stimulation. However, whereas sperm from 1.G2 and 6.F3L colonies did not seem to have any obvious issues, the sperm from 6.K2B colony, the one presenting the smallest sporosacs, presented an overall much lower sperm motility when compared to the sperm of an age-matched Cas9-only control colony (electronic supplementary material, video S1). This result suggests an association of GNL3 with the proper development of male gametes, the absence of which negatively affects their final motility. Alternatively, the lack of GNL3 protein in the mature sperm cells might directly affect their motility.

## 3. Discussion

To date, most GNL3 studies have been focused on vertebrate NS. In contrast, studies characterising invertebrate GNL3 have been scarce and sporadic across organisms, and generally not completely accounted for when inferring evolutionary hypotheses regarding how invertebrate GNL3 relates to NS and GNL3L in vertebrates.

We show that *GNL3* is expressed in *Hydractinia* i-cells and germline, and demonstrate its involvement in cell proliferation, animal growth, regeneration, and sperm motility. Our study establishes *GNL3* as a novel cnidarian stem cell and germline marker with deep ancestry, and opens new paths for a better understanding of stem cell systems and their evolution.

### 3.1. Invertebrate GNL3 paradigm shift

Based on the fact that human and mouse NS could not rescue the *GNL3* mutant phenotype in *S. pombe* and *C. elegans*, respectively [31, 33], but human GNL3L could rescue it in S*. pombe* [31], and on the involvement of invertebrate GNL3 in ribosome biogenesis [31–33], it has been proposed that vertebrate *GNL3L* is the homolog of invertebrate *GNL3* [17, 24]. We argue that this evidence is not enough to claim that invertebrate *GNL3* is more functionally related to *GNL3L* than it is to *NS*:

1. Rescue experiments with gene/protein expression from other organisms of distant phylogenetic position (i.e., xenorescues) do not always work, likely due to functional divergence or modification of binding sites occurring during sequence evolution.
2. In the *S. pombe* study [31], the expression of the closely-related *S. cerevisiae GNL3* (*nug1*) could not rescue the growth defect phenotype of *GNL3* (*Grn1*) mutants, while the distantly-related human *GNL3L* could partially restore it. This paradoxical result suggests a divergent evolution of the gene *GNL3* in this yeast species (supported by our phylogeny; figure 1C), making it appear to be more functionally similar to human *GNL3L* than to the closely-related *S. cerevisiae GNL3* and to human *NS*.
3. Interestingly, *D. melanogaster* and *C.elegans* GNL3 sequences do not cluster with GNL3L nor with NS protein families (electronic supplementary material, figure S1). This implies that *D. melanogaster* and *C. elegans GNL3* genes, while studied more in-depth [32, 33], have also diverged the most, potentially allowing for new functions to arise and others to disappear. Moreover, plant *GNL3* studies have not been considered when inferring the current evolutionary scenario. All this highlights the necessity of studying a variety of organisms from different phyla, since the focus on model organisms could potentially obscure the complete picture of the evolution and functions of a gene.

Our broad sampling for our phylogenetic domain analysis illustrates the presence of a GN3L_Grn1 domain at the N-terminus of all invertebrate GNL3 and vertebrate NS amino acid sequences, but the complete absence or a much lower GN3L_Grn1 domain sequence identity in vertebrate GNL3L sequences. Moreover, invertebrate GNL3 and NS sequences show a higher similarity in the presence and distribution of NLS sequences than they do to GNL3L. Overall, the most parsimonious evolutionary explanation is that invertebrate GNL3 and vertebrate NS share common ancestry, and that GNL3L arose secondarily as a novel gene during the vertebrate duplication event, becoming more evolutionary derived, contrary to what has been previously suggested [17, 24]. We thus propose a new paradigm for GNL3 evolution, where vertebrate GNL3L functionally diverged from its paralog, NS, as well as from its invertebrate ortholog, GNL3, while NS and invertebrate GNL3 share common ancestry and thus have retained more protein domain and functional identity (figure 6).

**Figure 6.**
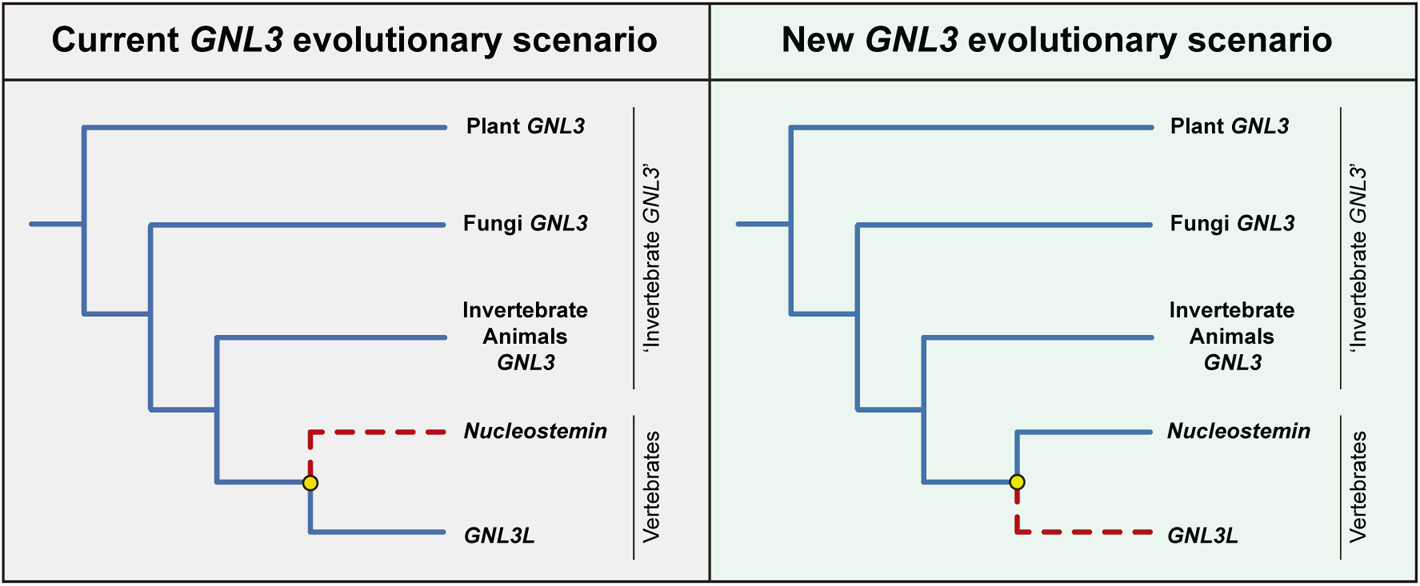
*GNL3* gene evolutionary scenarios. Left, current *GNL3* evolutionary scenario based on [17, 24]: Vertebrate *GNL3L* is hypothesized to be the direct descendant of invertebrate *GNL3*, and *Nucleostemin* arose as a novel gene with new functions following a duplication event during vertebrate evolution (yellow circle). Right, proposed new *GNL3* evolutionary scenario: Vertebrate *Nucleostemin* is the direct descendant of invertebrate *GNL3*, and *GNL3L* arose as a novel gene with new functions following a duplication event in the vertebrate clade (yellow circle).

This paradigm shift has important consequences on the interpretation of GNL3 functional evolution. Based on the new paradigm we propose, GNL3L could evolve novel functions acquired after the presumptive duplication event in the vertebrate clade, but also retain some ancestral functions that rely on the highly conserved domain MMR1_HSR1, some of which might be redundant with its paralog, NS. In contrast, NS and invertebrate GNL3 are more likely to share more ancestral functions, since both the GN3L_Grn1 and the MMR1_HSR1 domains have been conserved throughout evolution. In this manner, some GNL3L functions like the modulation of the transcriptional levels of oestrogen-related receptors [28], or the negative regulation of telomere length [27], are likely vertebrate GNL3L novelties, while vertebrate NS and invertebrate GNL3 involvement in other aspects such as regeneration, stemness, or genome stability might have been retained from the ancestral functions of GNL3. Further studies of invertebrate *GNL3* genes in different organisms will shed light on the evolution of GNL3, and on the paradigm shift in evolutionary scenarios we propose.

### 3.2. GNL3, growth, and regeneration

Experiments with *GNL3* mutants at the organismal level have demonstrated the indispensability of GNL3 for organism growth. For example, *C. elegans GNL3* knockouts display larval growth arrest [33], and *A. thaliana GNL3* mutants exhibit growth defects in both aerial and underground organs, leading to dwarf plants [36, 37]. Our results suggest that the slower growth rate observed in *Hydractinia GNL3* knockout colonies is due to reduced cell proliferation, similarly to what occurs in plants [37]. *Hydractinia GNL3* KO growth could be affected by a combination of other reasons in addition to reduced cell proliferation, however, such as impaired ribosomal subunit export leading to lower translation levels, or excessive cell differentiation; phenotypes observed in plant *GNL3* mutants [38] and in *NS* knockdowns of embryonic stem cells [19], respectively. Thus, further experimentation would be needed to specify the reason(s) for the slow growth phenotype observed in *Hydractinia GNL3* knockout colonies. A link could be made between *GNL3* / *NS* involvement in organismal growth ([33,36,37]; this study) and tumour growth [46, 47], highlighting the potential that *GNL3* / *NS* downregulation could have in developing cancer therapies to reduce tumour growth. Nanoparticle drug delivery systems targeting *NS* have already been tested with successful results, obtaining smaller-sized prostate cancer tumours [47].

The biological process of regeneration has long attracted researchers, and efforts have been made to broaden the therapeutic strategies in regenerative medicine. Studies *in vivo* on emerging model organisms with high regenerative capabilities, such as cnidarians and planarians, will help decipher the shared aspects of regeneration in multicellular organisms of different phylogenetic positions, highlighting features that are essential for regeneration [48, 49]. In *Hydractinia* and in planarians, head regeneration depends on cell proliferation, and the primary cellular source for blastema establishment is the migration of pluripotent stem cells (i-cells in *Hydractinia*, cNeoblasts in planarians) from the body to the prospective head [11,50,51]. Other strategies such as transdifferentiation, which occurs *in vitro* in the hydrozoan jellyfish *Podocoryne carnea* [52], or dedifferentiation, which takes place during newt lens regeneration [53] and during zebrafish heart and fin regeneration [54], have not yet been assessed during *Hydractinia* head regeneration, and thus their contribution cannot be discarded. The importance of GNL3 in regeneration processes has been illustrated by NS accumulation in dedifferentiating retinal pigmented epithelial cells during newt lens regeneration and in degenerating muscle fibres during newt limb regeneration [53], and by GNL3 requirement for complete head and tail regeneration in planarians [34]. Our results demonstrate the necessity of GNL3 for regeneration in *Hydractinia*, although the mechanism through which GNL3 functions during regeneration remains unknown. In combination with previous findings, our data suggests an ancient and evolutionarily-conserved role of GNL3 in regeneration, and infers that GNL3 might be essential for all regeneration processes, independently of the phylogenetic position of the regenerating organism, and of the cellular and molecular strategies used for regeneration. This emphasizes the relevance of GNL3 research for regenerative medicine progress.

### 3.3. GNL3 in sexual reproduction

Previous reports have illustrated the importance of GNL3 for sexual reproduction: NS was found present at high levels in mouse male germ cells, and its knockdown reduced germ stem cell proliferation potential *in vitro* [55]. Germline-specific *C.elegans GNL3* mutants were sterile, presenting defects in germline stem cell proliferation [33]. *NS* was differentially expressed in sterile *Xenopus* hybrid testes when compared to fertile non-hybrids [56]. In *A. thaliana*, *GNL3* mutants presented defects in reproductive fertility seen by the development of defective flowers, and the presence of unfertilised ovules [35, 36]. We showed *GNL3* expression in *Hydractinia* germ cells, oocytes, and proliferating spermatogonia. Moreover, one outcome of *Hydractinia GNL3* gene knockout we observed was the presence of defects in sexual polyps and the spawning of impaired sperm. Therefore, our results add to the body of evidence that *GNL3* disruption has negative effects on the germline and, more generally, on sexual reproduction, thus affecting the fitness of a species.

### 3.4. GNL3 cellular functions in different phylogenetic contexts

It has been shown that vertebrate NS perturbation reduces cell proliferation and can induce cell cycle arrest [45,57–60], and that plant *GNL3* mutants present inefficient cell proliferation and impaired cell cycle progression in their meristems [37], therefore affecting cell cycle dynamics. Our results show that *GNL3* disruption reduces the number of S-phase and M-phase cells without affecting the overall number of i-cells in *Hydractinia* larvae, strongly suggesting that cell cycle dynamics are affected. Interestingly, Qu and Bishop, 2012 [19] demonstrated that *NS* depletion in mouse embryonic stem cells reduced the number of S-phase cells due to a lengthened G1 phase of the cell cycle, which in turn led to increased differentiation. Based on our current data, we cannot pinpoint whether a similar mechanism governs the cell proliferation phenotype we obtained. Further experimentation would help determine the specific mechanisms involved.

Vertebrate *NS* depletion can induce apoptosis (i.e., programmed cell death) of different cancer and stem cell types [61–65] due to reduced cell viability. In *Hydractinia*, apoptosis occurs naturally as part of the metamorphosis process from larva to polyp, but only takes place sporadically in a few cells during larval growth and homeostasis [66]. Our results show that apoptosis does not take place when *GNL3* gene is knocked down in *Hydractinia* larvae, even while cell proliferation is affected. This suggests that *GNL3* is not crucial for *Hydractinia* i-cell viability, possibly thanks to the plasticity of cnidarian cells [5], which might allow them to overcome the insult and avoid apoptosis. Downregulation of vertebrate *NS* induces DNA damage, since it hampers the homology-directed repair of DNA damage foci that spontaneously occur during DNA replication of stem and progenitor cells [20]. NS also plays an important role in preventing telomere damage [21]. Hence, NS carries out essential functions in genome protection of actively-dividing cells. Our results show that *GNL3* knockdown does not induce spontaneous DNA damage in larval cells, but our approach does not discount the possibility that GNL3 might still be involved in DNA damage repair upon genotoxic stress. It has been proposed that the DNA damage repair function of GNL3 is a vertebrate NS innovation [17, 24]. This hypothesis contrasts with the results obtained in *A. thaliana*, however, where *GNL3* mutants presented higher sensitivity to treatments with genotoxic agents [37], suggesting plant GNL3 involvement in DNA damage repair processes. Similar experiments to those performed in plants and in *Hydractinia* could be performed in a wide range of organisms to shed light on the open question of whether the DNA damage repair ability of GNL3 is ancestral, and has been conserved throughout evolution, or whether it is a vertebrate novelty.

Several lines of work have demonstrated the involvement of NS [22–24], GNL3L [24], and invertebrate GNL3 [30–33,38] in different aspects of ribosome biogenesis, such as the biosynthesis of mature rRNAs or the cytoplasmic export and assembly of 60S ribosomal subunits. Our results strongly suggest that the knockdown of *GNL3* does not affect mature rRNA biosynthesis in *Hydractinia* (contrary to what was shown for GNL3L [24]), but do not exclude the possibility that other aspects of ribosome biogenesis, such as large subunit export and assembly, could be affected. Based on the fact that NS, GNL3L, and invertebrate GNL3 have been shown to be involved in one facet or another of ribosome biogenesis, it is likely that *Hydractinia* GNL3 also has some involvement in these processes. Further experimentation is needed to confirm this hypothesis.

Overall, invertebrate GNL3 and NS are seemingly involved in a myriad of cellular functions. These functions might vary depending on the organism, on the stem or tumour cell type, on the microenvironment surrounding *GNL3/NS*-expressing cells, and on particular cellular burdens such as genotoxic exposure or nucleolar stress. The fact that some interacting partners of NS, like p53 or mouse double minute 2 (MDM2) [14, 15], are missing in some organisms where GNL3 is present, increases even more the complexity of the functional evolution of this puzzling stem cell gene. Nonetheless, our results, in addition to our broad phylogenetic survey of GNL3 / NS functions, suggest that GNL3 has an ancient evolutionary-conserved function in stem cell regulation via the control of cell proliferation.

## 4. Opening Up

It has been proposed that stem cells are not a homologous entity shared among animals of different phylogenetic clades, and that stem cell-specific regulatory networks might have evolved independently in vertebrates and cnidarians [5]. The potential homology of stem cell types between different animal clades would be supported if more evidence of evolutionarily-conserved stem cell genes could be found regarding their presence and function. Thus, the identification of *GNL3* as a *Hydractinia* i-cell marker, and its presence in stem cells from both animals and plants, contrasts with the idea of stem cells evolving independently. Given the wide phylogenetic conservation of GNL3 proteins and their involvement in stem and germ cell regulation in plants and animals, we hypothesise that the ancestral *GNL3* gene might have been part of the original gene toolkit of the common ancestor of all eukaryotes. Our results, together with previous findings on *GNL3* in other organisms, suggest that the genetic control of stem cell regulation might present deep ancestry, and support a common evolutionary origin of stem cell types. More in-depth functional and mechanistic studies of the *GNL3* gene in a wide range of organisms would be invaluable to better understand its functions and the extent to which they have been conserved, shedding light into the biology and molecular basis of stem cell systems and how they have evolved.

## 5. Material and Methods

### 5.1. Animal husbandry and drug treatment

*Hydractinia* spawning, embryo and larvae culturing, larval metamorphosis induction, and adult colony breeding were performed as previously described [43]. A newly optimised Monday-Friday feeding regime was established, consisting exclusively of SEP-Art Artemia nauplii (INVE Aquaculture), which were consistently enriched two times with S.presso (SELCO) the day before colony feeding.

To induce DNA damage, larvae were incubated in the ribonucleotide reductase inhibitor hydroxyurea (HU; Sigma-Aldrich, St. Louis, MO. USA) at a concentration of 20mM in Millipore-Filtered Seawater (MFSW) for 3 hours.

### 5.2. YlqF/YawG GTPase family cluster map, GNL3 molecular phylogeny, and GNL3 domain analysis

To identify YlqF/YawG GTPase family genes in *Hydractinia*, tBLASTn searches were performed on *Hydractinia* genome and assembled transcriptomes, using human YlqF/YawG GTPase family protein sequences as bait. Protein sequence clustering was performed using CLANS2 [67], with a BLOSUM62 matrix and a *p*-value cutoff of 1e-1000. Sequences were retrieved from UniProtKB and by recovering the top 100 hits of BLAST searches while using each of *Hydractinia* YlqF/YawG GTPase family representatives as query. CD-HIT [68] was run with 99% identity to exclude sequence duplicates. The FASTA file containing the 1067 sequences used to obtain the cluster map (electronic supplementary material, figure S1) can be found in electronic supplementary material, file S1.

To perform GNL3 molecular phylogenetic analyses, a subset of GNL3 sequences found in electronic supplementary material, file S1 were handpicked to create a comprehensive list that encompasses plants, fungi, protists and most major animal clades (a number of GNL3 sequences not present in file S1 were retrieved from available online genomic and transcriptomic sources and added to the phylogenetic analyses; GNL2 sequences were also added to be used as outgroup; electronic supplementary material, file S2). A total of 112 full-length protein-coding sequences were aligned automatically using MAFFT v6.861b with the linsi options [69]. The final alignment file is provided in electronic supplementary material, file S3. ProtTest 3 [70] which calls PhyML for estimating model parameters [71] was used to select the best-fit model of protein evolution for the alignment, which was LG + I + gamma + F (“LG” indicates the substitution matrix, “I” specifies a proportion of invariant sites, “gamma” specifies gamma-distributed rates across sites, and “F” specifies the use of empirical amino acid frequencies in the dataset). Maximum likelihood (ML) analyses were performed using RaxML v 8.2.9 [72]. ML branch support was estimated using non-parametric bootstrapping (500 replicates). The resulting tree was rooted with the GNL2 clade in FigTree v 1.4.4 (http://tree.bio.ed.ac.uk/software/figtree/) and compiled in Adobe Illustrator.

To analyse the different GNL3, GNL3L and NS protein domains, we used the respective amino acid sequences from a broad subset of organisms belonging to each clade of the phylogenetic analyses shown in Figure 1C as query. Domain analyses were performed using PFAM (https://pfam.xfam.org/) and Motif Scan (https://myhits.sib.swiss/cgi-bin/motif_scan).

### 5.3. *In situ* hybridisation

For colorimetric *in situ* hybridisation, all samples were first relaxed in in 4% MgCl_2_ 1:1 MFSW-mqH_2_O for 20 minutes, and then fixed in Fix 1 (4% PFA + 0.2% glutaraldehyde + 0.1% Tween-20 in MFSW) for 90 seconds at room temperature. This was followed by removal of Fix 1 and incubation in ice-cold Fix 2 (4% PFA + 0.1% Tween-20in MFSW) for 90 minutes at 4°C. After fixation, three washes of 15 minutes in ice-cold PBS containing 0.1% Tween-20 (PTw) were performed. For permeabilization and storage, samples were dehydrated in increasing concentrations of methanol in PTw (25%, 50%, 75%, 100%), and stored at −20°C for at least 24h. Samples were rehydrated by decreasing concentrations of methanol in PTw (75%, 50%, 25%) and washed three times in PTw. Samples were then placed in a heat block at 85°C for 20 minutes to inactivate endogenous alkaline phosphatases. This step was followed by 10-minute washes with 1% Triethanolamine in PTw, then with 6μl/ml and 12μl/ml acetic anhydride diluted in 1% Triethanolamine-PTw. After several washes in PTw, samples were transferred into a 24-well plate and pre-hybridised in hybridisation buffer (4M urea, 5x SSC pH 7.0, 1% SDS, 0.1% Tween-20, 100 ug/ml tRNA, and 50 ug/ml heparin in DEPC-treated mqH_2_O) without probes for 2-5 hours at 55°C. After pre-hybridisation, Digoxigenin-labelled antisense RNA probe for *GNL3* was preheated at 90°C for 10 minutes and added to fresh hybridisation buffer immediately before incubation with the samples at 55°C for 36-60 hours. Following hybridisation, samples were washed with decreasing concentrations of hybridisation buffer in 2x SSC (at 55°C), followed by decreasing concentrations of 0.2X SSC in PTw (at room temperature). After post-hybridisation washes, two 10-minute washes in maleic acid buffer (-MAB-100mM Maleic acid, 150mM NaCl, pH 7.5) containing 0.1% Triton X-100 (MABT) followed. Samples were then blocked in blocking solution (1/10 Roche Blocking Buffer - ref. 11096176001-in MAB) for at least one hour at room temperature, followed by antibody incubation (Anti-Digoxigenin-AP, Fab fragments; Sigma-Aldrich) at 1:5000 dilution in blocking solution, overnight at 4°C. The following day, samples were washed six times in MABT and then incubated in alkaline phosphatase (AP) buffer (100 mM NaCl, 50 mM MgCl_2_, 100 mM Tris −pH 9.5, and 0.5% Tween-20 in mqH_2_O) containing 0.33 mg/ml NBT and 0.165 mg/ml BCIP. This solution was refreshed every 1-2 hours during the first day of development and twice a day the following days. When desired, the reaction was stopped by washing samples several times in PTw. Samples were mounted in TDE (97% 2,2′-Thiodiethanol and 3% 1x PBS) before microscopy.

For larvae double fluorescent *in situ* hybridisation, samples were fixed, stored and treated as for colorimetric ISH unless noted otherwise. Hybridisation was performed using a Digoxigenin-labelled probe for *GNL3* and a Fluorescein-labelled probe for *Piwi1* or *PCNA*. Following post-hybridisation washes, endogenous peroxidase activity was quenched by incubating samples twice for 30 minutes in 3% H_2_O_2_ at room temperature. Samples were first incubated overnight at 4°C with an antibody Anti-Digoxigenin-POD, Fab fragments (Sigma-Aldrich) at 1:1500 dilution in blocking solution. Samples were washed six times in MABT, then washed in PTw for 45 minutes, and incubated in tyramide development solution (2% dextran sulfate, 0.0015% H_2_O_2_, 0.2 mg/ml iodophenol, and 1x Alexa Fluor 594 Tyramide reagent - B40957-in PTw) for 10 minutes at room temperature. After five 15-minute washes in PTw, fluorescence was quenched with 100mM glycine solution (pH 2.0) for 10 minutes at room temperature and washed again four times in PTw. Overnight incubation at 4°C with an antibody Anti-Fluorescein-POD, Fab fragments (Sigma-Aldrich) at 1:1500 dilution in blocking solution was followed by washes in MABT, PTw and tyramide development solution as before (containing 1x Alexa Fluor 488 Tyramide reagent (B40953) in place of Alexa Fluor 594). Nuclei were stained using Hoechst 33342 and samples were mounted in Fluoromount (Sigma-Aldrich) before confocal imaging analysis. Primers used for probe synthesis can be found in electronic supplementary material, file S4.

### 5.4. Immunofluorescence, TUNEL and EdU experiments

For immunofluorescence, larvae were relaxed, fixed and stored as for ISH, using PBS + 0.1% Tween-20 (PTw) instead of MFSW + 0.1% Tween-20 in the fixation buffers. Samples were rehydrated by decreasing concentrations of methanol in PTw (75%, 50%, 25%) and four 15-min washes PTx (0.02% Triton X-100 in 1x PBS) at room temperature followed. Samples were then permeabilised with 0.3% Triton X-100 in 1x PBS for 20 minutes at room temperature and washed again three times with PTx prior to a blocking step of 3-5 hours in 0.2 μm-filtered blocking solution (10% Bovine Serum Albumin −BSA-, 5% Normal Goat Serum - NGS-in PBS1X). After blocking, samples were incubated overnight at 4°C in primary antibody anti-PH3 (anti-Histone H3 phospho-Ser10; Arigo Biolaboratories), anti-GH2A.X (Anti-phospho-Histone H2A.X (Ser139) Antibody, clone JBW301; EMD Millipore), or anti-Piwi1 [10] diluted 1:150, 1:200, or 1:100, respectively, in blocking solution. The following day, samples were washed four times for 10 minutes each in PTx + 5% BSA, followed by two long (1h) washes in PTx + 5% BSA. Samples were then blocked again in fresh blocking solution for 1-3 hours at room temperature, then incubated for 1h at room temperature with secondary antibody Goat-anti-Rabbit 568 (Invitrogen) or Goat-anti-Mouse 488 (Invitrogen) at 1:500 dilution in blocking solution. Four washes of 30 minutes in PTx + 5% BSA followed, and samples were left washing in PTx + 5% BSA overnight at 4°C. The following day, samples were rinsed in PTx prior to nuclei staining.

For TUNEL assays, samples were relaxed as detailed above and fixed for 3 hours at room temperature in TUNEL Fix (0.1M Hepes, 0.05M EGTA, 0.01M MgSO_4_, 0.02% Triton X-100, and 4% PFA in MFSW), followed by three 15-minute washes in PTw. Samples were dehydrated, stored, rehydrated, permeabilised, washed, and blocked as for immunofluorescence, then rinsed in 1x PBS. For staining of apoptotic cells, we used the In Situ Cell Death Detection Kit Fluorescein (Millipore Sigma, 11684795910) following manufacturer’s recommendations. For positive controls, samples were incubated at 37°C for 20 minutes in 2 units of DNAseI diluted in 50 μl of 1x DNase buffer (from RNAqueous-Micro Total RNA Isolation Kit; Ambion), prior to TUNEL enzyme reaction.

To detect cells in S-phase of the cell cycle, EdU (Life Technologies C10340, Carlsbad, CA, USA) was added to solutions containing samples of interest, to a final concentration of 150 µM for different lengths of time, depending on sample type (5 minutes for 2 dpf and 3 dpf larvae; 10 minutes for primary polyps; 15 minutes for 8 dpf and 21 dpf larvae; 20 minutes for adult feeding and sexual polyps). Samples were then fixed, dehydrated, stored, rehydrated, permeabilised, washed, and blocked as for immunofluorescence. The Click-iT EdU detection reaction was carried out for 1 hour at room temperature following manufacturer’s recommendations. When combined with immunofluorescence or fluorescent *in situ* hybridisation, the Click-iT EdU detection reaction was performed at the end of the protocol, prior to nuclei staining.

In all cases, nuclei were stained using Hoechst 33342 and samples were mounted in Fluoromount (Sigma-Aldrich).

### 5.5. shRNA design, synthesis and electroporation

Design and synthesis of shRNAs targeting *GNL3* and a scrambled-*GNL3* control, as well as the electroporation procedure and survivorship assessment, were performed as previously described [43], with the exception that all electroporations were executed with a single shRNA at a concentration of 1500 ng/μl. Forward and reverse oligonucleotides of 66 bases in length that correspond to the DNA templates for shRNA *in vitro* transcription can be found in electronic supplementary material, file S5.

### 5.6. RT-qPCR and rRNA assessment

Larval RNA extraction, cDNA synthesis and qPCR analyses were performed as previously described [43]. Results were normalised to *Eef1alpha* housekeeping gene expression. Relative transcript expression levels of *GNL3* KDs were obtained using the delta-delta-ct method relative to scrambled shRNA controls. These relative expression levels are depicted as arbitrary units (A.U.) in figure 3. Three or more independent biological replicates were performed per experiment. Primer sequences are found in electronic supplementary material, file S5.

For rRNA assessment, larval RNA extraction was performed as previously described [43]. rRNA quality was checked using the Agilent 2100 Bioanalyzer. Results regarding rRNA levels and ratios are shown in electronic supplementary material, figure S7.

### 5.7. Generation of CRISPR-Cas9 mutant colonies

The three CRISPR sgRNAs targeting *GNL3* were designed using CRISPRscan, ordered from Synthego and resuspended in TE buffer (Synthego) at a concentration of 100-150 μM. sgRNAs were aliquoted and kept at −20°C until use. We avoided off-target matches by scanning the *Hydractinia* genome assembly at http://crispor.tefor.net. The sequences of the sgRNAs used in this study are found in in electronic supplementary material, file S6. Cas9 protein (CP02; PNA BIO) was reconstituted in nuclease-free water to a concentration of 30 μM, aliquoted and kept at −80°C until use. Microinjection mixtures consisted of Cas9 protein at 6 μM, a combination of the three sgRNAs at approximately 5 μM concentration each, 100mM KCl, and Dextran (Alexa Fluor 555; Invitrogen) at 1mg/ml concentration, all diluted in nuclease-free water. For Cas9-only controls, TE buffer (Synthego) was added instead of sgRNAs. Before microinjection, mixtures were incubated for 10 minutes at room temperature, then centrifuged at 14,000 rpm for 10 minutes at room temperature. Within 80 minutes of spawning and following microinjection, eggs were fertilised and incubated at 28°C for 3 hours to enhance Cas9 protein activity during early stages of development, and then transferred to 18°C for the rest of the experiment. Injected embryos were cultured for 3 days in MFSW until larval metamorphosis was induced as previously described [43]. Only 4-5 metamorphosing larvae were added per slide, to allow colony growth assessment without a crowding effect and to avoid the fusion of neighbouring colonies. Slides with established primary polyps were transferred to aquariums to avoid algal growth and enhance stolonal expansion. Primary polyps were mouth-fed daily with smashed brine shrimp until colonies were large enough to be fed with whole nauplii. If two colonies of any experimental condition came into contact or fused, these were excluded from the analyses. Once colony growth data was collected (figure 4), a single colony on each slide was maintained as a founder for further experimentation.

### 5.8. Knockout genotyping

Following metamorphosis of individual larvae and colony growth analyses, 1-2 polyps per colony were taken for genomic DNA (gDNA) extraction. gDNA extraction buffer (0.01M Tris pH8.0, 0.05M KCl, 0.3% Tween-20, 0.3% NP40, 0.001M EDTA, 0.5mg/mL Proteinase K; [73]) was freshy prepared and placed on ice. Feeding polyps were dissected from the colony and placed inside the lid of a 0.5 mL PCR tube, where as much seawater as possible was removed, before 20 µL of extraction buffer was pipetted on to the polyp. Tubes were centrifuged briefly before being placed at 55°C for 2-3 hours, and vortexed every 30-60 mins during incubation. Following incubation at 55°C, Proteinase K was inactivated by incubation at 98°C for 5 mins. PCR was conducted using 5 µL or 10 µL of gDNA as input template with Takara ExTaq DNA polymerase (RR001A) and *GNL3* specific primers that flanked the 3 predicted cut sites (electronic supplementary material, figure S8, S10; file s6). Resulting fragments were analysed via agarose gel electrophoresis.

Three colonies (1.G2, 6.K2B and 6.F3L) were chosen for further analysis of CRISPR/Cas9-induced mutations. Following PCR and gel electrophoresis, bands of interest were excised and purified using the QIAquick Gel Extraction kit (Qiagen, Cat. #28704) and ligated into the pGEM-T Vector System (Promega, Cat. #A1360). Chemically competent DH5-alpha *E. coli* bacteria (ThermoFisher, EC0111) were transformed and plated on LB-agar plates containing ampicillin (100µg/mL), IPTG (0.5mM) and X-gal (80µg/mL). Individual colonies were picked and verified to contain insert before being grown in overnight cultures of LB-broth containing ampicillin (100µg/ml). Plasmid DNA was extracted using the Qiagen Miniprep Kit. Plasmid clones derived from individual bacterial colonies were Sanger-sequenced by Psomagen (https://psomagen.com/) using vector primers and analysed in Geneious to identify mutations.

### 5.9. Imaging, cell counting and stolonal area quantifications

Individual polyp and colony images were acquired with a digital camera (Zeiss Axiocam ERc 5s) attached to a stereo microscope (Zeiss Stemi 508). Images of specimens from ISH experiments were taken with a digital camera (Zeiss Axiocam HRc) attached to a compound light microscope (Zeiss Imager.M2). Sperm motility videos were acquired with a Rolera EM-C^2^ high-speed camera (QImaging) attached to a compound light microscope (Zeiss Imager.M2). Following immunofluorescence, EdU labelling or fluorescent ISH, animals were imaged using a confocal microscope (Zeiss LSM 710). When comparisons between animals within an experiment were required the same scanning parameters were used for all conditions of each independent experiment. All maximum intensity projections of z-stacks were generated using Fiji [74].

For EdU^+^, PH3^+^, and Piwi1^+^ cell counting, larvae were compressed between the slide and cover slip to enable imaging of the full larval depth, and confocal z-stacks of approximately 10-15μm were used. EdU^+^ cells were highlighted using custom thresholding and counted using 3D Object Counter in Fiji. PH3^+^ and Piwi1^+^ cells were manually counted in Fiji. For quantification of colony stolonal surface in *GNL3* KO experiments, light microscopy images were used. The animals’ perimeters were outlined and areas were quantified using Fiji.

### 5.10. Graphs and statistics

Box plots in figure 3 and electronic supplementary material, figures S4, S8 and S10 were generated using BoxPlotR [75]. The remaining graphs were designed in Excel and compiled using Adobe Illustrator.

For assessment of RT-qPCR statistical significance, we utilised the delta-Ct values, and performed Shapiro-Wilk tests to check for normality, followed by two-tailed Student’s T-Tests. For evaluation of 28s / 18s rRNA ratio statistical significance, we first performed Shapiro-Wilk tests to check for normality, followed by two-tailed Student’s T-Tests. For statistics related to cell counting, normality was tested using Shapiro-Wilk tests, and Mann-Whitney U nonparametric tests were performed for two-way comparisons. Fisher’s exact tests were chosen to analyse results of figure 4B based on 2 x 2 contingency tables. The statistical significance of stolonal area and polyp number comparisons between control and *GNL3* KO polyp colonies was determined by two-way ANOVA tests followed by post hoc Bonferroni corrections (electronic supplementary material, file S7). Statistical significance for all quantitative comparisons is indicated (***) where p<0.01 and (*) where p<0.05. Two-way comparison tests and Fisher’s exact tests were conducted at http://www.socscistatistics.com, two-way ANOVA tests at http://www.statskingdom.com, and post hoc tests at http://www.graphpad.com.

## Author’s contributions

G.Q.A.: conceptualization, formal analysis, investigation, validation, visualization, writing—original draft, writing—review and editing; D.J.: investigation, formal analysis, validation, visualization, writing—review and editing; C.S.: conceptualization, funding acquisition, project administration, resources, supervision, validation, writing—review and editing. All authors gave final approval for publication and agreed to be held accountable for the work performed therein.

## Data accessibility

Accession numbers and source databases from the protein sequences used in figure 1C are given in electronic supplementary material, table S1.

## Supporting information

Sup_FileS1 (Cluster map FASTA)

Sup_FileS2_phylogeny_input_fasta_112

Sup_FileS3_phylogeny_alignment_final_112

Sup File S4 (primers for riboprobes)

Sup File S5 (shRNA and RT-qPCR primers)

Sup File S6 (sgRNAs and Genotyping primers)

Sup File S7 (2-Way ANOVA and Bonferroni corrections)

## Acknowledgements

We thank Dr. Uri Frank for sending us an aliquot of the Piwi1 antibody, Dr. Mark Martindale for sharing microinjection and microscopy equipment, Dr. Leonardo Ibarra Castro for helping to improve our *Hydractinia* and brine shrimp culture system, and Maddison Harman for the *Hydractinia* photo from figure 1.

