## Supplementary_Information for "*GNL3* is an evolutionarily-conserved stem cell gene influencing cell proliferation, animal growth, and regeneration in the hydrozoan *Hydractinia*"

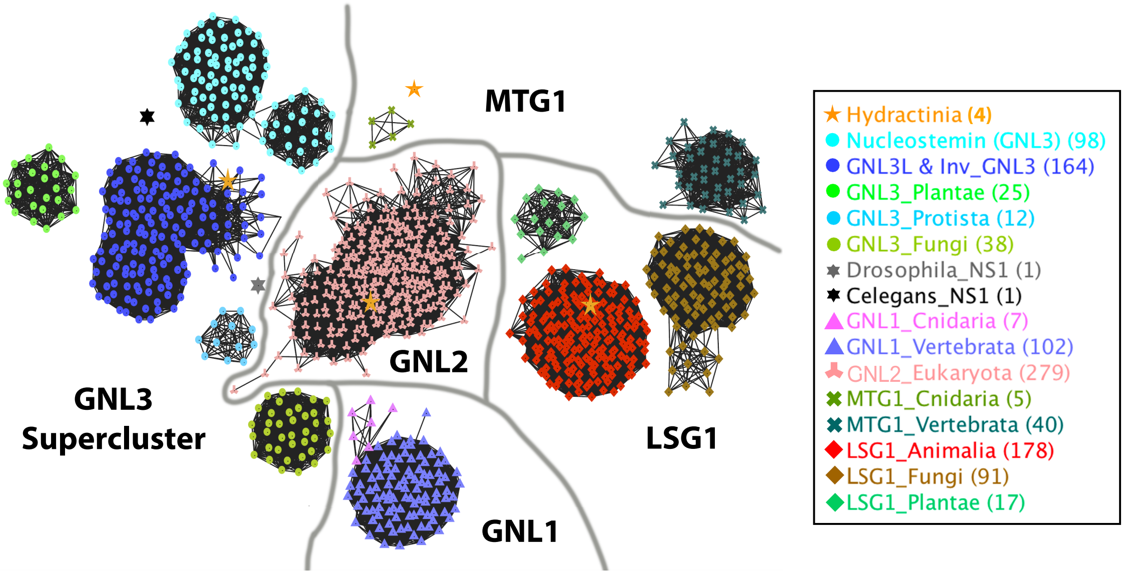


**Figure S1. YlqF/YawG GTPase family protein cluster map.** Sequence-similarity-based clustering of protein sequences from all major YlqF/YawG GTPase family groups (GNL1, GNL2, GNL3, MTG1, LSG1). BLASTP p-value <1e-1000. Each symbol represents a protein sequence (see legend on the right - numbers in parentheses represent the number of sequences in each protein cluster). The 1067 sequences encompass all four eukaryotic kingdoms (electronic supplementary material, file S1). The four *Hydractinia* sequences are depicted by orange stars. *D. melanogaster* (*Drosophila*) and *C.elegans* GNL3 protein sequences are depicted by a grey and a black star, respectively. Note how their GNL3 sequences do not have a direct connection with any of the GNL3 clusters, suggesting that their GNL3 amino acid sequences are more derived than most other animals. Inv = invertebrate. All connections with p-value <1e-1000 are shown.

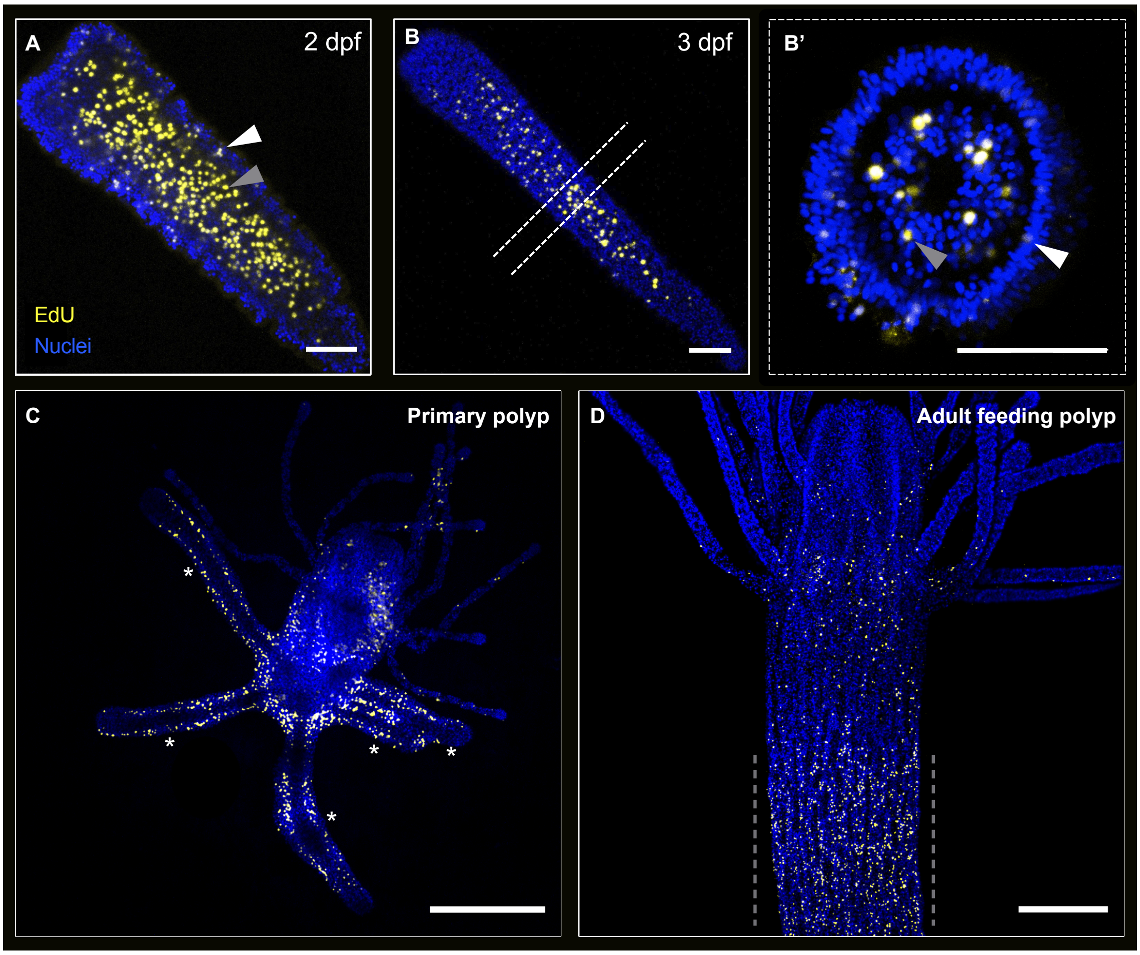


**Figure S2. Sites of cell proliferation in *Hydractinia* larvae and polyps.** (A) 2 dpf larva showing cells that have incorporated EdU (yellow) mostly in the larval endoderm (grey arrowhead) and a few cells in the larval ectoderm (white arrowhead). (B) 3 dpf larva with $EdU^+$ cells mostly present in the larval endoderm. (B') Transverse section of the mid-region of a 3 dpf larva, as shown in (B) by dashed lines, presenting strong EdU incorporation in endodermal cells, and weak EdU signal in some ectodermal cells. Arrowheads as in (A). (C) Aboral view of a primary polyp revealing EdU incorporation in cells of the polyp body column, in some of the tentacles, and in the stolons (asterisks). (D) Adult feeding polyp displaying EdU incorporation in cells of the body column and tentacles. Note the high concentration of $EdU^+$ cells in the i-cell band-like area at the base of the polyp (defined by grey dashed lines). All nuclei are stained in blue. All images shown were projected from confocal stacks. dpf = days post-fertilisation. Scale bars: 50 μm in A-B'; 200 μm in C-D.

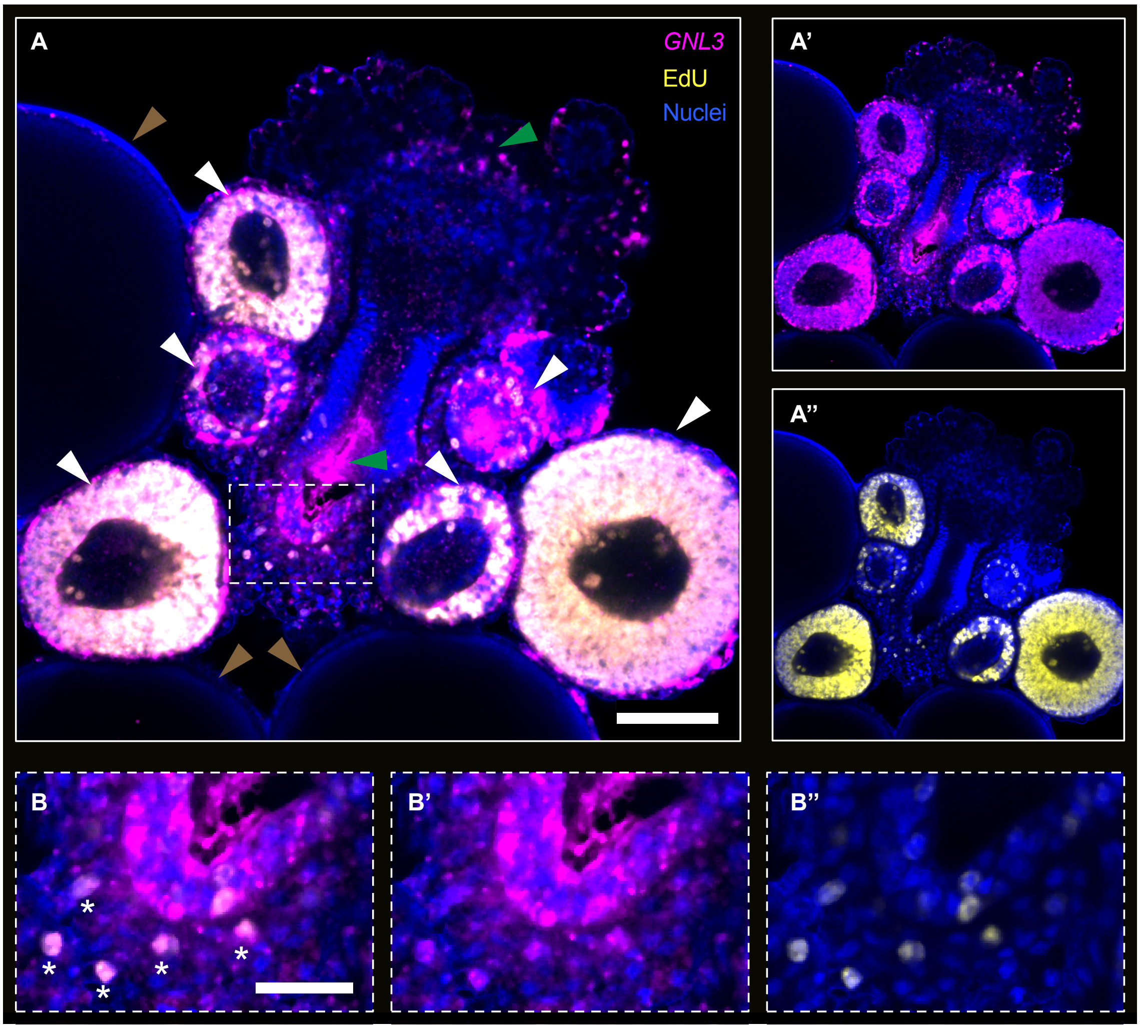


**Figure S3. *GNL3* is expressed in the germinal zone in male sexual polyps and co-localizes with EdU in spermatogonia.** (A) Fluorescent ISH of *GNL3* (magenta) co-stained with EdU (yellow) showing co-localisation in spermatogonia of small and middle-size sporosacs (white arrowheads), but the absence of both markers in large sporosacs (brown arrowheads) of a male sexual polyp. *GNL3* is also expressed in the germinal zone as well as in some cells surrounding it (defined by a white rectangle). Green arrowheads point to non-specific *GNL3* riboprobe staining. (A'-A'') Same image as in (A) showing *GNL3* and EdU individual channels, respectively. (B) Detail of the region outlined in (A) displaying strong *GNL3* expression in the germinal zone in contrast with the absence of EdU incorporation in most germ cells. Some cells neighbouring the germinal zone (white asterisks) contain both markers, and could potentially be germ cells migrating from the germinal zone towards nearby sporosacs. (B'-B'') Same image as in (B) showing *GNL3* and EdU individual channels, respectively. All nuclei are in blue. Images shown correspond to a single confocal plane. Scale bars: 50 μm in A; 20 μm in B.

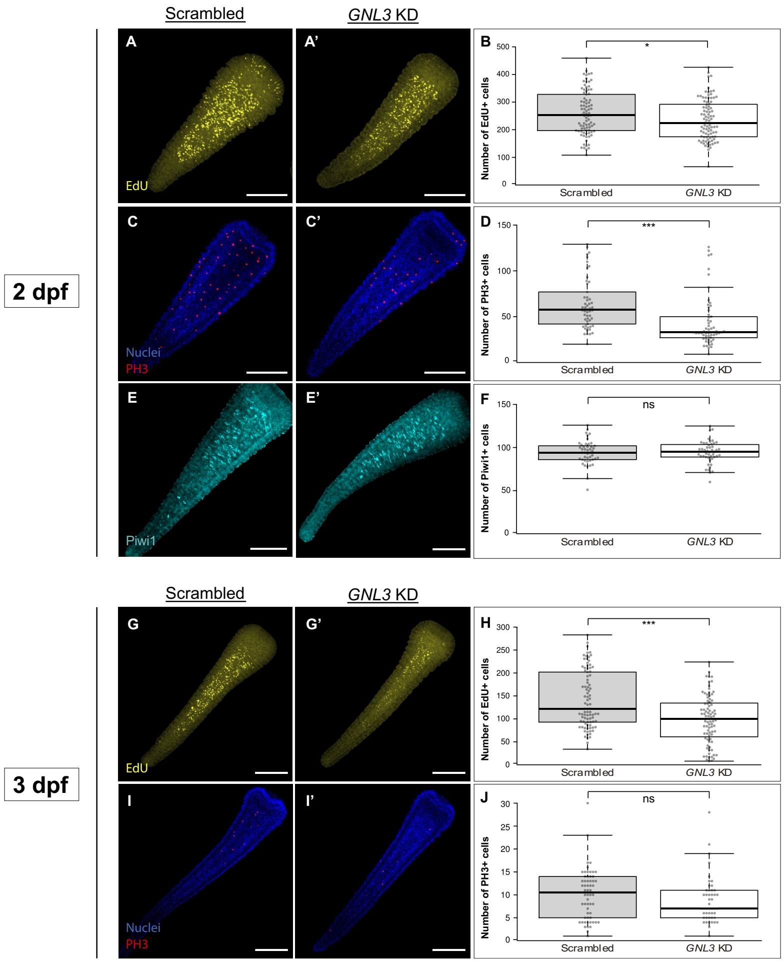


**Figure S4. *GNL3* knockdown significantly reduces the number of EdU$^{+}$ and PH3$^{+}$ cells without affecting the number of Piwi1$^{+}$ cells in larvae of different ages**. (A-A', C-C', E-E', G-G', I-I') Representative images of 2 dpf or 3 dpf larvae showing EdU$^{+}$ cells (yellow, A-A', G-G'), PH3$^{+}$ cells (red, C-C', I-I'), or Piwi1$^{+}$ cells (E-E') in scrambled and *GNL3* KD larvae. In C-C' and I-I' nuclei are stained in blue. (B, D, F) Box plot showing the number of EdU$^{+}$, PH3$^{+}$ or Piwi1$^{+}$ cells in 2dpf scrambled and *GNL3* KD larvae. (H, J) Box plot showing the number of EdU$^{+}$ ad PH3$^{+}$ cells in 3dpf scrambled and *GNL3* KD larvae. Centre lines show the medians; box limits indicate the 25th and 75th percentiles (first and third quartiles); whiskers extend 1.5 times the interquartile range from the 25th and 75th percentiles; each quantified sample is represented by a grey circle. In all cases, the full depth of the larvae was imaged and images shown were projected from confocal stacks. EdU quantifications were combined from 3 independent experiments (2dpf scrambled, n = 86; *GNL3* KD, n = 91; 3dpf scrambled, n = 89; *GNL3* KD, n = 83), whereas PH3 and Piwi1 quantifications were combined from 2 independent experiments (2dpf scrambled, n = 54; *GNL3* KD, n = 58 for PH3; 3dpf scrambled, n = 58; *GNL3* KD, n = 47 for PH3; 2dpf scrambled n = 45; *GNL3* KD n = 47 for Piwi1). dpf = days post-fertilisation. ns = non-significant; * = p-value≤0.05; *** = p-value≤0.01. All scale bars: 100 µm.

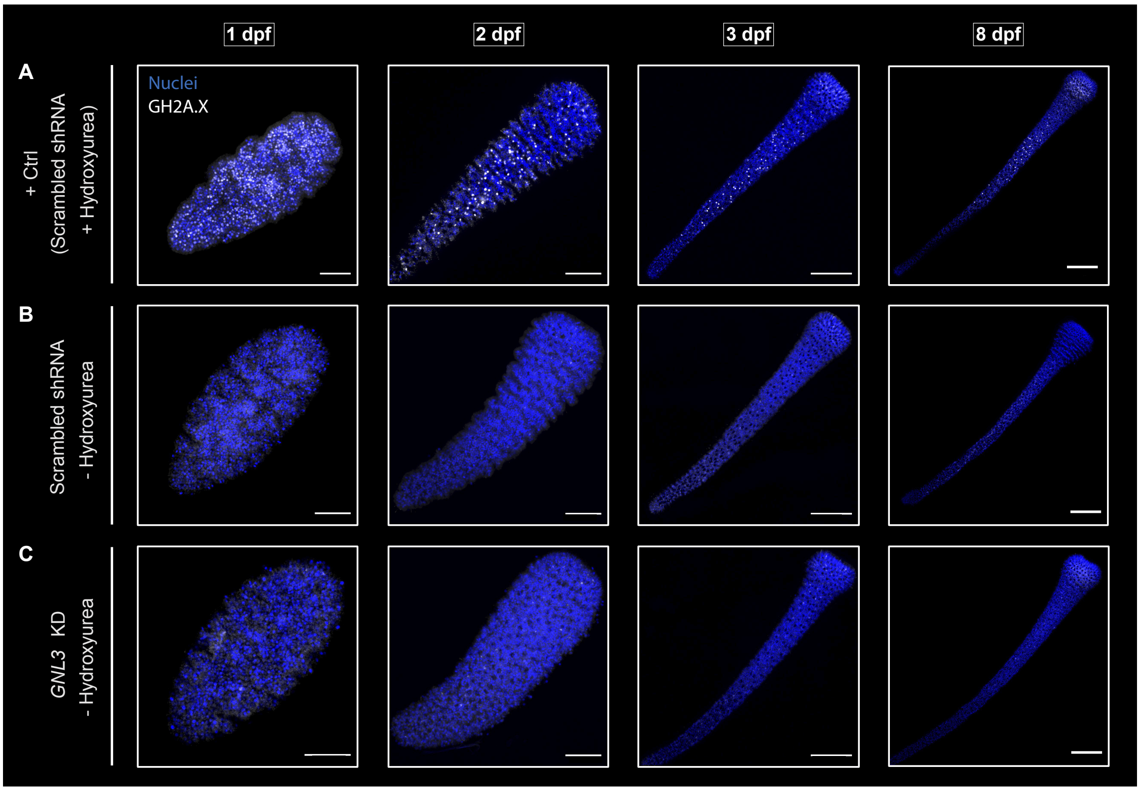


**Figure S5. *GNL3* knockdown does not generate spontaneous DNA damage in larvae**. (A) Representative larval images at different timepoints following hydroxyurea (HU) treatment to induce DNA damage prior to fixation (+ Ctrl). GH2A.X$^{+}$ cells (white) represent those cells in which HU has induced DNA damage. The pattern of GH2A.X$^{+}$ cells is reminiscent of that of proliferating cells (see electronic supplementary material, figure S2). (B) Representative larval images at different timepoints following electroporation of scrambled shRNA control, not treated with hydroxyurea. Note the almost complete absence of GH2A.X labelling. (C) Representative larval images at different timepoints following *GNL3* knockdown, not treated with hydroxyurea. The almost complete absence of GH2A.X labelling indicates that knockdown of *GNL3* does not induce spontaneous DNA damage. Nuclei are stained in blue. In all cases, n ≥ 30. All images shown were projected from confocal stacks. dpf = days post-fertilisation. Scale bars: 50 μm in 1 dpf and 2 dpf; 100 μm in 3 dpf and 8 dpf.

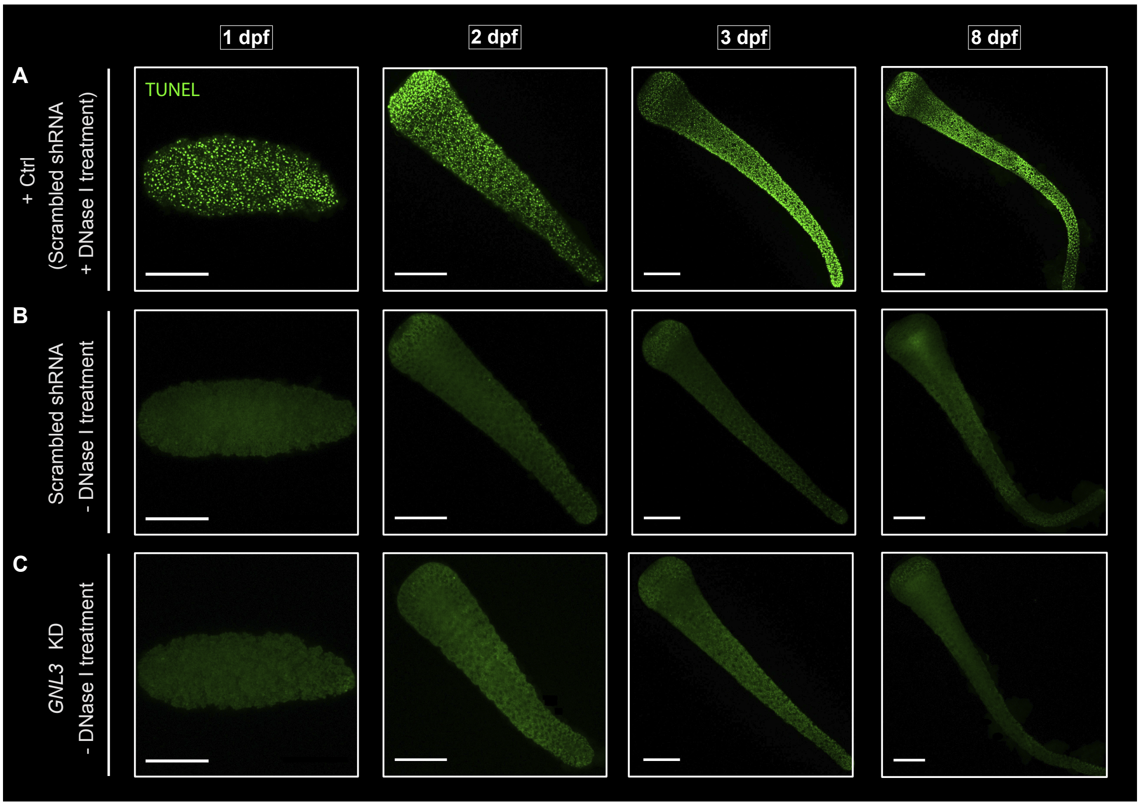


**Figure S6. *GNL3* knockdown does not induce apoptosis in larvae**. (A) Representative larval images at different timepoints in the positive control (+ Ctrl), where fixed larvae were treated with DNase I to induce DNA strand breaks prior to developing the TUNEL enzyme reaction. Most cells were labelled by TUNEL (green). (B) Representative larval images at different timepoints following electroporation of scrambled shRNA, not treated with DNase I. Note the absence of TUNEL labelling. (C) Representative larval images at different timepoints following *GNL3* knockdown, not treated with DNase I. The absence of TUNEL labelling indicates that the knockdown of *GNL3* does not induce apoptotic cell death. In all cases, n ≥ 30. All images shown were projected from confocal stacks. dpf = days post-fertilisation. All scale bars: 100 μm.

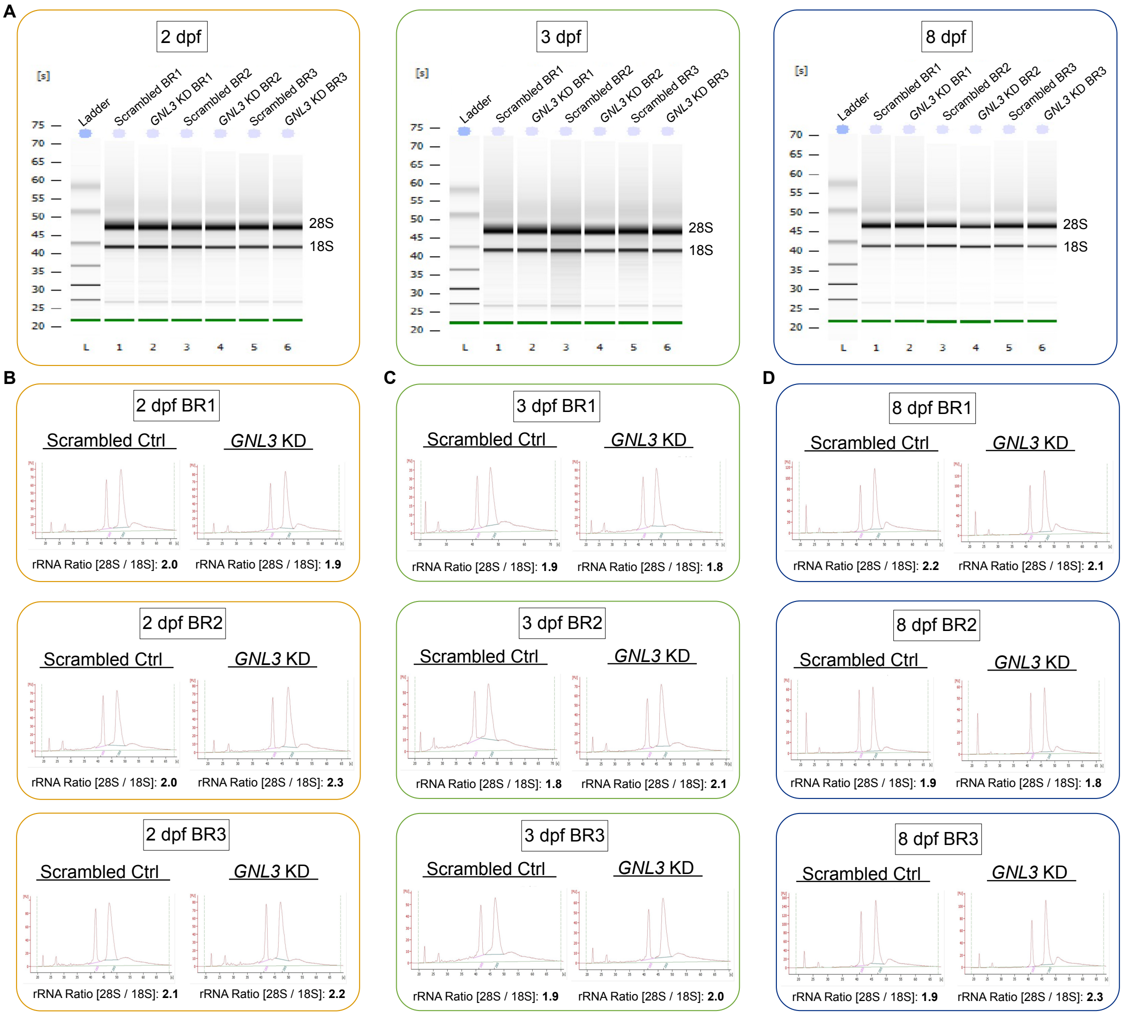


**Figure S7. *GNL3* knockdown does not affect 28S and 18S rRNA levels nor 28S / 18S rRNA ratios**. (A) Simulated gel electrophoresis images obtained from the Bioanalyzer after running three independent biological replicates (BR) of 2 dpf, 3 dpf, and 8 dpf larval RNA samples from scrambled and *GNL3* KD conditions. (B) Bioanalyzer electropherograms for 2 dpf larval RNA samples extracted from scrambled control and *GNL3* knockdown samples. Three independent biological replicates (BR) are shown. 28S / 18S rRNA ratios given by the Bioanalyzer are shown at the bottom of each electropherogram. (C) Same as (B) but for 3 dpf larval RNA samples. (D) Same as (B) but for 8 dpf larval RNA samples. Overall, GNL3 knockdown yields equivalent 18S and 28S rRNA levels, as well as comparable 28S / 18S rRNA ratios to scrambled controls. rRNA = ribosomal RNA. dpf = days post-fertilisation.

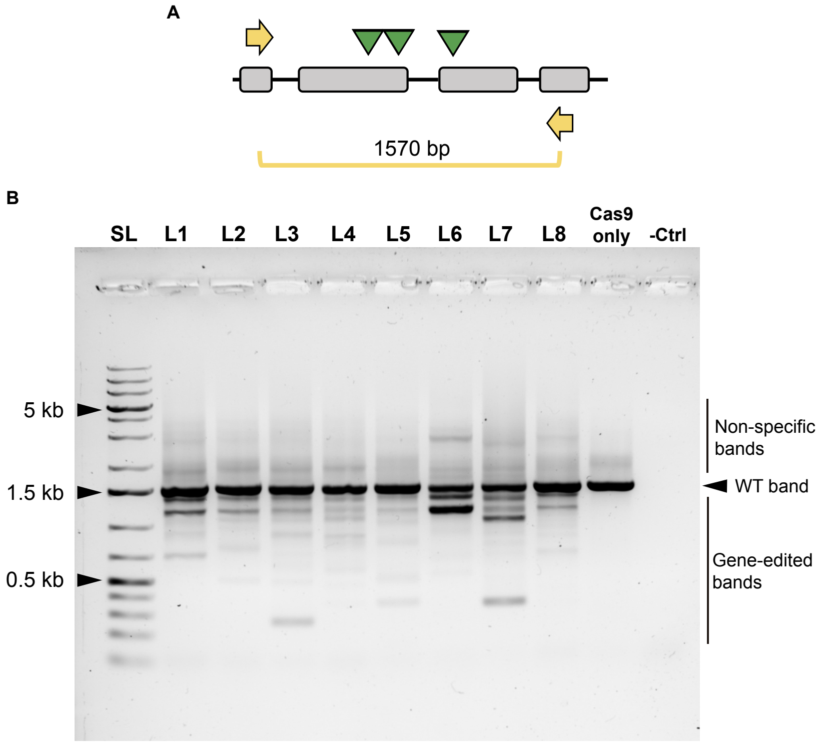


**Figure S8. CRISPR/Cas9-mediated knockout of *GNL3* induces mosaic gene editing in *Hydractinia* larvae**. (A) Schematic depicting exons four to seven of *Hydractinia GNL3* gene, focused on the region where CRISPR/Cas9 gene editing was targeted and genotyping primers designed. Exons (grey boxes), introns (black lines), target sites of sgRNAs (green triangles), genotyping primers (yellow arrows). 1570 bp represents the length of the sequence amplified by the primers. (B) Agarose gel electrophoresis image showing amplicons of *GNL3*, using the primers indicated in (A), from 8 individual 1 dpf larvae resulting from embryos injected with sgRNA/Cas9 complexes (L1-L8). A single larva injected with Cas9 protein only is shown as a control (Cas9 only), alongside a negative (no template) control (-Ctrl). SL = size ladder. 0.5kb, 1.5kb, and 5kb bands of the ladder are indicated. The wild-type band corresponds to the unedited 1570 bp product amplified from the *GNL3* gene (WT). Non-specific bands that are larger than 1.5kb in length were always found when amplifying this region of *Hydractinia GNL3* gene, possibly due to the presence of highly repetitive sequences promoting the formation of secondary and tertiary structures that migrated slower in the gel than the linear amplicon. In L1-L8 samples we cannot discard the possibility that some bands in this region of the gel are due to CRISPR/Cas9-induced insertions. Amplicons shorter than 1.5kb are present in sgRNA/Cas9-microinjected larval samples, but not in the Cas9-only control, and indicate the presence of deletions induced by CRISPR/Cas9 gene editing. Note that all edited larvae are mosaic as indicated by multiple PCR amplicons representing distinct gene editing events. bp = base pairs; dpf = days post-fertilisation.

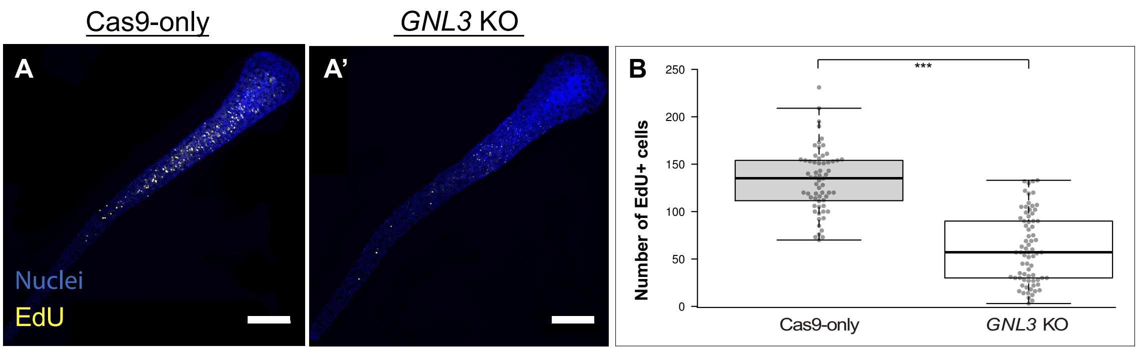


**Figure S9. *GNL3* knockout reduces number of EdU$^+$ cells in 21 dpf larvae**. (A-A') Representative images of 21 dpf larvae showing EdU$^+$ cells (yellow) for Cas9-only and *GNL3* KO conditions. Nuclei are stained in blue. (B) Box plot showing the number of EdU$^+$ cells for Cas9-only and *GNL3* KO conditions. Centre lines show the medians; box limits indicate the 25th and 75th percentiles (first and third quartiles); whiskers extend 1.5 times the interquartile range from the 25th and 75th percentiles; each quantified sample is represented by a grey circle. EdU quantifications were combined from 2 independent experiments. For Cas9-only, n = 63; for *GNL3* KO, n = 73. In all cases, the full depth of the larvae was imaged and images shown were projected from confocal stacks. *** = p-value≤0.01. dpf = days post-fertilisation. Scale bars: 100 μm.

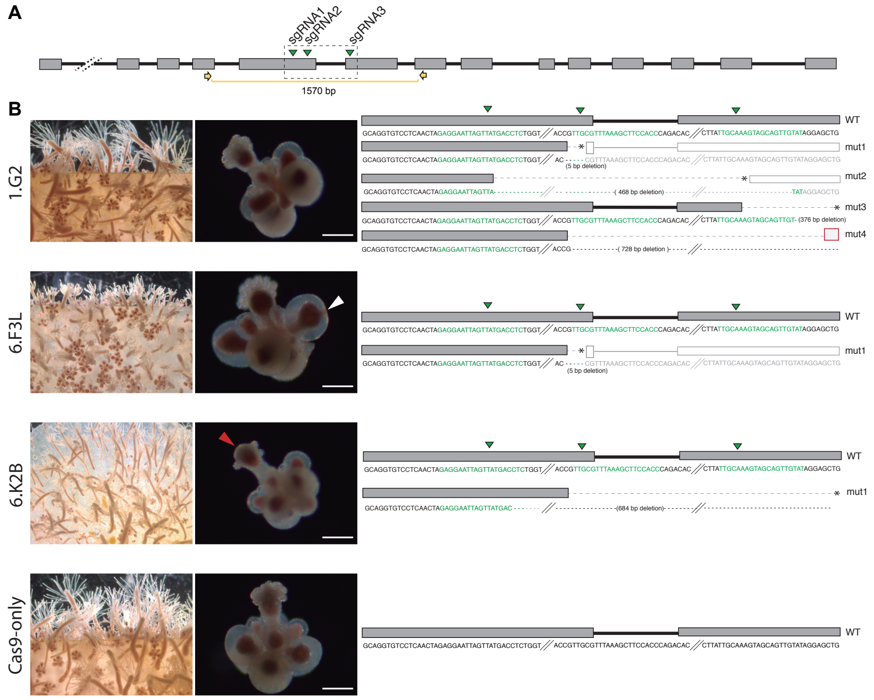


**Figure S10. Characterisation of CRISPR/Cas9-induced *GNL3* mutants.** (A) Schematic depicting the intron-exon structure of the *GNL3* locus. Exons (grey boxes), introns (black lines), target sites of sgRNAs (green triangles), genotyping primers (yellow arrows). 1570 bp represents the length of the sequence amplified by the primers. Box with dashed lines represents the area depicted on the far right in (B). (B) *GNL3* KO colonies derived from microinjection of embryos with sgRNA/Cas9 (1.G2, 6.F3L, and 6.K2B) alongside a colony derived from a Cas9-only injected embryo (Cas9-only). An image of each F0 colony (left), a single sexual polyp from each colony (middle), and mutations revealed by PCR, cloning and Sanger sequencing of *GNL3* amplicons (right) are shown. 6.F3L sexual polyps showed defects in sporosac structure (white arrowhead), while 6.K2B sexual polyps exhibited a smaller oral region (red arrowhead) and smaller sporosacs compared to age-matched controls. To the right of each colony are schematics of *GNL3* gene structure with exons (filled grey boxes), introns (black lines), and target sites of sgRNAs (green triangles) shown. Mutations resulting in non-sense mutations (*) and missense mutations (grey box with red outline) are shown. Nucleotide sequences of each mutation are shown beneath each schematic, with sgRNA target sequences shown in green. Generated mutants all maintained a wild type allele (WT), and each mutant also had different variations in *GNL3* editing resulting from different-sized deletions between sgRNA1 and sgRNA3. Colony 1.G2 had four detected *GNL3* mutations (mut1 – mut4), 6.F3L had a single detected mutation (mut1), and 6.K2B also had a single detected *GNL3* mutation (mut1). bp = base pairs. Scale bars: 200 µm.

### Supplementary Files Legends:

Supplementary Video S1. Video showing sperm motility from a Cas9-only male colony (left) compared to sperm motility from a *GNL3* knockout male colony (6.K2B colony; right). Note that while most Cas9-only sperm moves extremely fast, most *GNL3* knockout sperm moves remarkably slow. All sperm was collected within 5 minutes from the spawning event, and sperm motility was recorded for 3 minutes. The video was assembled in Fiji at 8 fps (frames per second).

Supplementary Table S1. Table listing the full species names and accession numbers of all the protein sequences used to create the GNL3 tree in figure 1C as well as the abbreviated names used in the tree. The source database or website for all sequences is also indicated.

Supplementary File S1. FASTA file containing all sequences used for electronic supplementary material, figure S1.

Supplementary File S2. FASTA file including all sequences used for the phylogenetic tree shown in figure 1C.

Supplementary File S3. Alignment of all sequences used for the phylogenetic tree shown in figure 1C.

Supplementary File S4. List of the primers used for riboprobe synthesis.

Supplementary File S5. List of the shRNA oligonucleotide sequences used for *in vitro* transcription of shRNAs, and of the RT-qPCR primer sequences used in this study.

Supplementary File S6. Sequences of the three sgRNAs used in this study, which were co-injected with Cas9 protein to create *GNL3* CRISPR/Cas9-mediated knockout mutants. Primers used for genotyping larvae and polyps are also listed.

Supplementary File S7. Detailed information of the Two-Way ANOVA statistical analyses and post-hoc Bonferroni corrections performed for figure 4D-E.
